# The chromatin structuring protein HMGA2 influences human subtelomere stability and cancer chemosensitivity

**DOI:** 10.1101/544320

**Authors:** Syed Moiz Ahmed, Priya Dharshana Ramani, Stephen Wong Qi Rong, Xiaodan Zhao, Roland Ivanyi-Nagy, Tang Choong Leong, Clarinda Chua, Zhizhong Li, Hannes Hentze, Iain Tan Bee Huat, Jie Yan, Ramanuj DasGupta, Peter Dröge

## Abstract

The transient build-up of DNA supercoiling during the translocation of replication forks threatens genome stability and is controlled by DNA topoisomerases (TOPs). This crucial process has been exploited with TOP poisons for cancer chemotherapy. However, pinpointing cellular determinants of the best clinical response to TOP poisons still remains enigmatic. Here, we present an integrated approach and demonstrate that endogenous and exogenous expression of the oncofetal high-mobility group AT-hook 2 (HMGA2) protein exhibited broad protection against the formation of hydroxyurea-induced DNA breaks in various cancer cells, thus corroborating our previously proposed model in which HMGA2 functions as a replication fork chaperone that forms a protective DNA scaffold at or close to stalled replication forks. We now further demonstrate that high levels of HMGA2 also protected cancer cells against DNA breaks triggered by the clinically important TOP1 poison irinotecan. This protection is most likely due to the recently identified DNA supercoil constraining function of HMGA2 in combination with exclusion of TOP1 from binding to supercoiled substrate DNA. In contrast, low to moderate HMGA2 protein levels surprisingly potentiated the formation of irinotecan-induced genotoxic covalent TOP1-DNA cleavage complexes. Our data from cell-based and several *in vitro* assays indicate that, mechanistically, this potentiating role involves enhanced drug-target interactions mediated by HMGA2 in ternary complexes with supercoiled DNA. Subtelomeric regions were found to be extraordinarily vulnerable to these genotoxic challenges induced by TOP1 poisoning, pointing at strong DNA topological barriers located at human telomeres. These findings were corroborated by an increased irinotecan sensitivity of patient-derived xenografts of colorectal cancers exhibiting low to moderate HMGA2 levels. Collectively, we uncovered a therapeutically important control mechanism of transient changes in chromosomal DNA topology that ultimately leads to enhanced human subtelomere stability.

**Author Summary:** DNA replication fork stability in rapidly dividing cancer cells is of utmost importance for the maintenance of genome stability and cancer cell viability. Cancer cells efficiently prevent fork collapse into lethal double strand breaks as a first line of defense during replication stress, but the corresponding protective mechanisms often remain elusive.

Uncontrolled high levels of DNA supercoiling that are generally regulated by topoisomerases can cause replication stress and are major threats to fork stability. Using a multidisciplinary approach, we identified a possible regulatory mechanism of replication stress, which appears to involve mitigating the consequences of DNA topological changes by the oncofetal replication fork chaperone HMGA2.

Our work provides mechanistic insights into the control of DNA damage triggered by clinically important anti-cancer drugs, which is mediated by the replication fork chaperone HMGA2. We thereby also identify HMGA2 expression as a predictive therapeutic marker, which could allow clinicians to take informed decisions to prevent tumor recurrence and improve survival.

## Introduction

DNA transactions in all life forms require that protein complexes translocate at high speed along the two intertwined strands of the DNA double helix. If these complexes are not free to rotate around the DNA template, the translocation process will induce changes in local DNA topology. In this context, the introduction of positive (+) and negative (−) DNA supercoiling are the most profound consequences, and the generation of such transient supercoil waves in the chromatin appears to be inevitable (1, 2). However, under certain conditions, their uncontrolled build-up can also become a threat to genome stability (3).

High levels of (+) DNA supercoiling generated during replication are particularly dangerous to genomes. The free energy stored in topologically overwound DNA in front of replicative helicases can stall the translocation of replisomes. It has frequently been argued that (+) supercoiling could also contribute to unscheduled replication fork reversal (4–9), which would result in pathological chicken foot fork structures. Both scenarios, i.e. extensive genome-wide fork stalling with or without fork regression, can ultimately contribute to fork collapse into lethal DNA double strand breaks (DSBs) (4–11). Owing to the highly dynamic nature of DNA topological changes during replication, however, these processes are experimentally very challenging to study and thus represent an often overlooked or unappreciated threat to genome stability. This contrasts, for example, with the much better understood protein-mediated fork regression processes that are critical for stalled fork stability, repair and restart (12, 13).

Two classes of enzymes, type 1 and type 2 topoisomerases (TOP1/2), have evolved to maintain supercoiling homeostasis during replication by supercoil relaxation via transient DNA strand breaks. The great biological importance of these reactions is highlighted by the fact that treatment of cancer cells with clinically successful TOP1/2 poisons results in the trapping of these enzymes in covalent TOP-DNA complexes (3, 14, 15). The resulting impairment of supercoil relaxation in the parental DNA during replication and the impediments to replisomes can trigger fork stalling, precatenane formation, uncontrolled fork regression and, ultimately, fork collapse into lethal DSBs (3, 7, 8, 10, 12, 15–21). In this context, it has recently been argued that TOP1-trapping by camptothecin also leads to stress due to the build-up of DNA supercoiling in yeast (22). A second, more established scenario is that the trapping of TOP1 in covalent complexes with DNA (TOP1cc) downstream of replisomes by, for example, the camptothecin drug irinotecan, leads to replication fork run-off that converts single-stranded DNA lesions into highly cytotoxic DSBs (23, 24). Taken together, TOP1/2 poisons, through their various modes of actions during replication and other DNA transactions, have been successfully exploited for chemotherapy of common human cancers. However, identifying cellular determinants of the best clinical response to TOP poisons remains a challenge (17).

The human oncofetal, non-histone chromatin factor high-mobility group AT-hook 2 (HMGA2) is evolutionarily highly conserved in mammals. *HMGA2* is expressed during early developmental stages and is aberrantly re-activated in many cancers (25, 26). We recently reported that HMGA2 exhibits replication fork chaperone activity in human and the heterologous yeast and *Escherichia coli* systems, where HMGA2 significantly reduced both fork collapse into cytotoxic DSBs and the formation of chromosomal aberrations. The protein is found in close proximity to stressed replication forks, and we presented evidence that HMGA2 might, at least in part, act there through interfering with fork regression (27). However, the mechanistic details underlying this novel fork chaperone function of HMGA2 remained to be uncovered.

In this context, subsequent *in vitro* studies provided some interesting leads. Single DNA manipulation experiments showed that HMGA2 binds with high affinity to both (+) and (−) supercoiled DNA (+/− scDNA) and readily forms unique HMGA2-scDNA complexes via a novel mode of intramolecular DNA segment-bridging. This form of supercoil scrunching, as well as the replication fork chaperone function, requires the presence of AT-hook DNA-binding domains of HMGA2 (27, 28). Furthermore, our recent unbiased high-throughput compound screen in combination with biochemical assays pointed at a functional interaction between HMGA2 and human TOP1 (29). We therefore hypothesized that HMGA2, in addition to or in conjunction with its proposed scaffold forming function at HU-induced stalled forks, may be a cancer/stem cell-specific modulator of dynamic changes in chromatin structure involving DNA supercoiling that contributes to replication fork stability during replication stress.

In this study, we tested this hypothesis and demonstrate using a combination of biochemical, single DNA manipulation, and cell-based assays that HMGA2 has an important cellular function in dealing with DNA topological challenges which occur during replication stress triggered by the clinically important TOP1 poison irinotecan. In particular, human subtelomeres contain narrow regions of enhanced vulnerability to replication stress. Our integrated approach also included patient-derived xenografts of colorectal cancers and corroborated the role of HMGA2 as an important cancer cell-specific determinant of the efficacy of irinotecan. Hence, besides uncovering an important control mechanism of localized chromosomal DNA supercoiling, our study also suggests that future personalized therapeutic interventions should consider HMGA2 protein expression levels in human tumors.

## Results

### HMGA2 expression broadly protects cancer cells against hydroxyurea-induced replication stress

Chemotherapeutic agents that target replicative DNA polymerases, such as aphidicolin or hydroxyurea (HU), induce replication stress through fork stalling. Extended HU exposure for up to 24h results in replication fork-associated DSBs, which are distinct from apoptotic DNA responses to drug treatment (30–32). The resulting genomic DNA fragments vary in size and amount, and can be taken as a measure of the extent of fork collapse induced by these agents.

We have recently shown that HMGA2 can contribute to the protection of stalled forks by reducing DSB formation in HU-challenged cells. This fork chaperone function was ascribed to the formation of protective scaffolds that HMGA2 forms through DNA binding at stalled forks (27), but the precise chaperoning mechanism remained elusive. In the previous study, we utilized two HT1080 clonal human cancer cell lines and human embryonic stem cells (hESCs) that permitted us to control endogenous HMGA2 expression levels via sh and si RNA, respectively (27). We have extended these investigations here and asked whether this fork chaperone function is active in other cancer cell types. In addition to the HT1080 fibrosarcoma cells, we utilized cervical carcinoma HeLa and lung adenocarcinoma A549 cancer cell models in which exogenous human *HMGA2* is expressed at different levels and can be compared to the corresponding parental cells, which do not express detectable protein levels as determined by Western blot analysis (**Fig. 1**) (HeLa parental and HeLa P2/P8/P19; A549 parental and A549 1.3/1.5/1.6 cell lines; (37)). We also employed human non-small lung carcinoma H1299 cells in which expression of endogenous *HMGA2* had been knocked out (KO) by the CRISPR-Cas9 technology (**Fig. 1**) (H1299 parental and HMGA2 KO cells).

**Fig 1.**
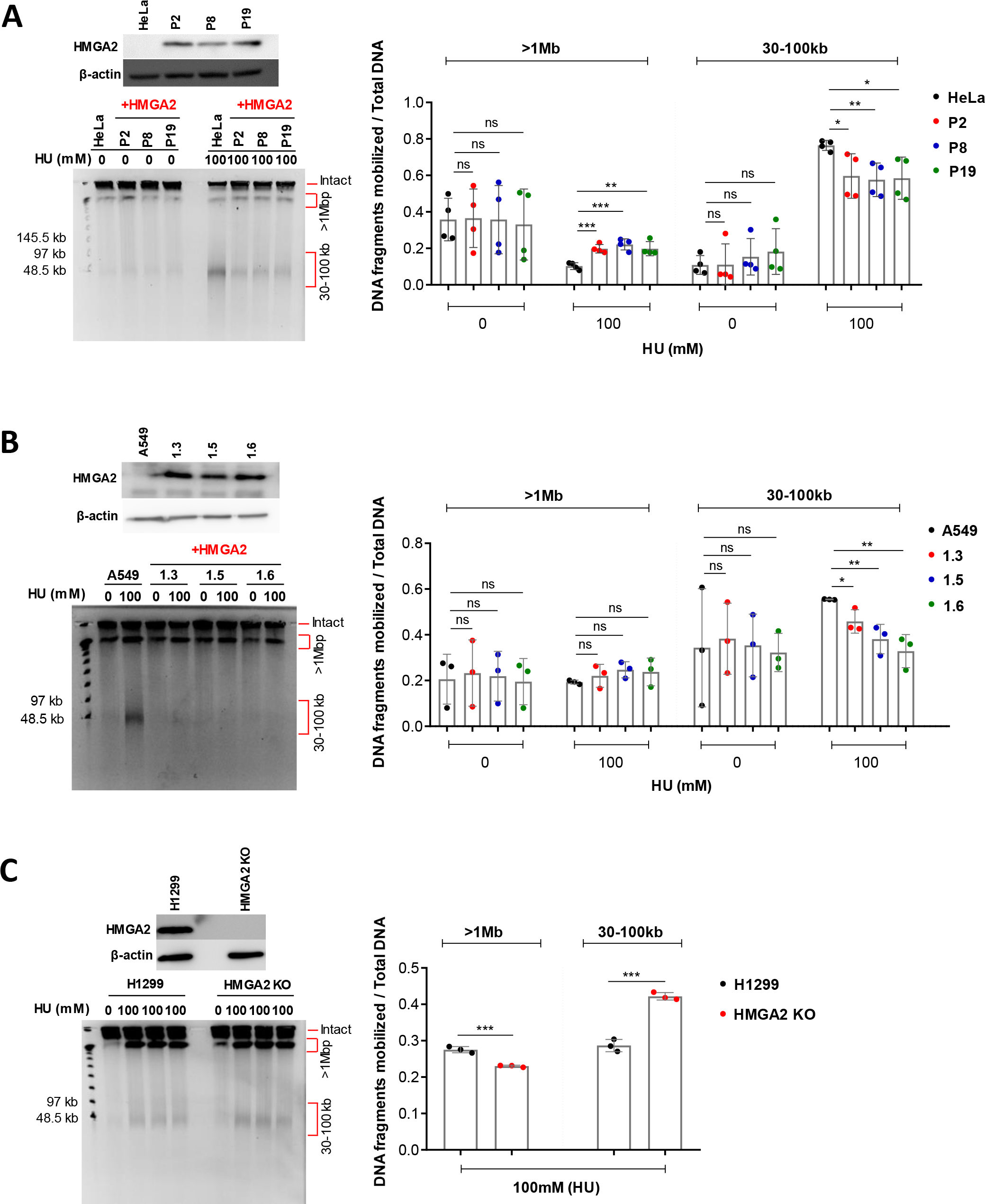
HMGA2 protects against HU-induced DSBs. **(A)** A western blot shows corresponding HMGA2 expression levels in HeLa cells (parental and three recombinant human HMGA2-expressing clonal HeLa cell lines) (top left). Representative PFGE analysis of DSB formation in HeLa cells in response to 24h incubation with HU (left). Quantification of HU-induced DNA fragments (>1Mb and 30-100kb fractions) (right) was done by ImageJ software with each fragment fraction normalized to total DNA loaded (n=4 independent experiments). Error bars show s.d. Unpaired two-tailed t-tests. ns not significant, * p < 0.05, ** p < 0.01. **(B)** A western blot shows corresponding HMGA2 expression levels in A549 cells (parental and three recombinant human HMGA2-expressing cell lines) (top left). Representative PFGE analysis of DSB formation in A549 cells in response to 24h incubation with HU (left). Quantification of HU-induced DNA fragments (>1Mb and 30-100kb fractions) (right) was done by ImageJ software with each fragment fraction normalized to total DNA loaded (n=3 independent experiments). Error bars show s.d. Unpaired two-tailed t-tests. * p < 0.05, ** p < 0.01, *** p < 0.001, **** p < 0.0001. **(C)** A western blot shows corresponding HMGA2 expression levels in H1299 cells (parental and HMGA2 KO) (top left). Representative PFGE analysis of DSB formation in H1299 cells in response to 24h incubation with HU (left). Quantification of HU-induced DNA fragments (>1Mb and 30-100kb fractions) (right) was done by ImageJ software with each fragment fraction normalized to total DNA loaded (n=3 independent experiments). Error bars show s.d. Unpaired two-tailed t-tests. * p < 0.05, ** p < 0.01, *** p < 0.001, **** p < 0.0001.

The extent of DSB formation that results from HU-induced replication stress was investigated through pulsed field gel electrophoresis (PFGE). This type of assay is based on the analysis of cell populations without DNA extraction/purification and can provide a quantitative measure of the corresponding genomic DNA fragments, which can be up to several megabase pairs (Mbp) in length. Larger chromosomal fragments cannot be mobilized and remain trapped in the loaded cell plugs. In agreement with our previous study (27), we found that 24h HU treatment generated two distinct high molecular weight genomic DNA fragment fractions (30-100kb and >1Mbp) (**Fig. 1 and S1 Fig.**). For each sample, the DNA fragment fractions were normalized to the total DNA (total DNA = two distinct DNA fragment fractions + the intact genomic DNA fraction in the well that is too large to be mobilized by the electric fields). Strikingly, HMGA2 protected against HU-induced DSB formation across all cell lines (including our previously employed HT1080 C1 and C2 cells, which were used here as controls (27) (**S1 Fig.**)). Notably, the most consistent reduction in DSB formation was seen in the 30-100kb DNA fragment fraction (**Fig. 1 and S1 Fig.**). Hence, we conclude that HMGA2 appears to have a general role as fork chaperone across various types of HU-treated cancer cells.

In a recent independent study that involved single-molecule analysis, we found that HMGA2 binds to plectonemic supercoils with very high affinity, subsequently leading to their constrainment (28). This raised the possibility that HMGA2 could have a function in the resolution of DNA topological problems that may arise during replication stress in front of replication forks. In order to probe further into this hypothesis, we employed the chemotherapeutic drug irinotecan, which specifically targets human TOP1.

### HMGA2 expression determines DSB formation during SN38-induced replication stress

Human TOP1 in collaboration with TOP2 plays a key role in maintaining supercoiling homeostasis in the parental DNA during replication fork translocation (3, 20). Hence, the trapping of TOP1 in covalent DNA complexes by the metabolically active form of irinotecan, termed SN38, has two major consequences: first, the more established mode of action is through TOP1cc formation, which triggers replication fork run-off events that convert single-stranded DNA lesions into highly cytotoxic DSBs (23, 24). The second, still debated consequence is the build-up of excessive (+) supercoiling in the parental DNA, which either promotes fork stalling/regression and collapse into DSBs, or the formation of pre-catenanes behind the fork which can only be resolved by the action of TOP2 (15, 16, 18).

In order to test our hypothesis that HMGA2 is a modulator of transient chromosomal supercoiling generated during DNA replication, we used SN38 to treat the four cancer cell line models introduced above. We found that 48h drug exposure also generated the two-distinct high molecular weight genomic DNA fragment fractions seen with HU-treated cells (30-100kb and >1Mbp) (**Fig 2**). Quantification of the fragment fractions and normalization to the total DNA revealed that HMGA2 expression efficiently protected H1299 and A549 cells against SN38-induced DSBs, most notably in the 30-100kb fragment fraction (**Fig 2A and 2B**). To further validate our results, we measured caspase activation and found that HMGA2 levels negatively correlated with both the extent of SN38-induced DSBs and apoptosis (**Fig 3A and 3B**), and, as expected, positively correlated with cell survival (**Fig 3A**; **right panel**). Surprisingly, however, HMGA2 expression in HeLa and HT1080 cell models clearly had the opposite effect, i.e. HMGA2 sensitized cells to the genotoxic effects of SN38 (**Fig 2C and 2D; also see S2 Fig**). Determination of caspase activation over a range of drug concentrations revealed that HMGA2 expression directly correlated with increased SN38-induced DSBs and apoptosis (**Fig 3C and 3D**) and negatively correlated to cell survival (**Fig 3C**; **bottom panel**). Furthermore, the generation of the 30-100kb fragment fraction in HeLa cells and the increase in the amount of these fragments due to HMGA2 expression in HeLa P2 cells was already detectable after 6h SN38 treatment (**S2 Fig**).

**Fig 2.**
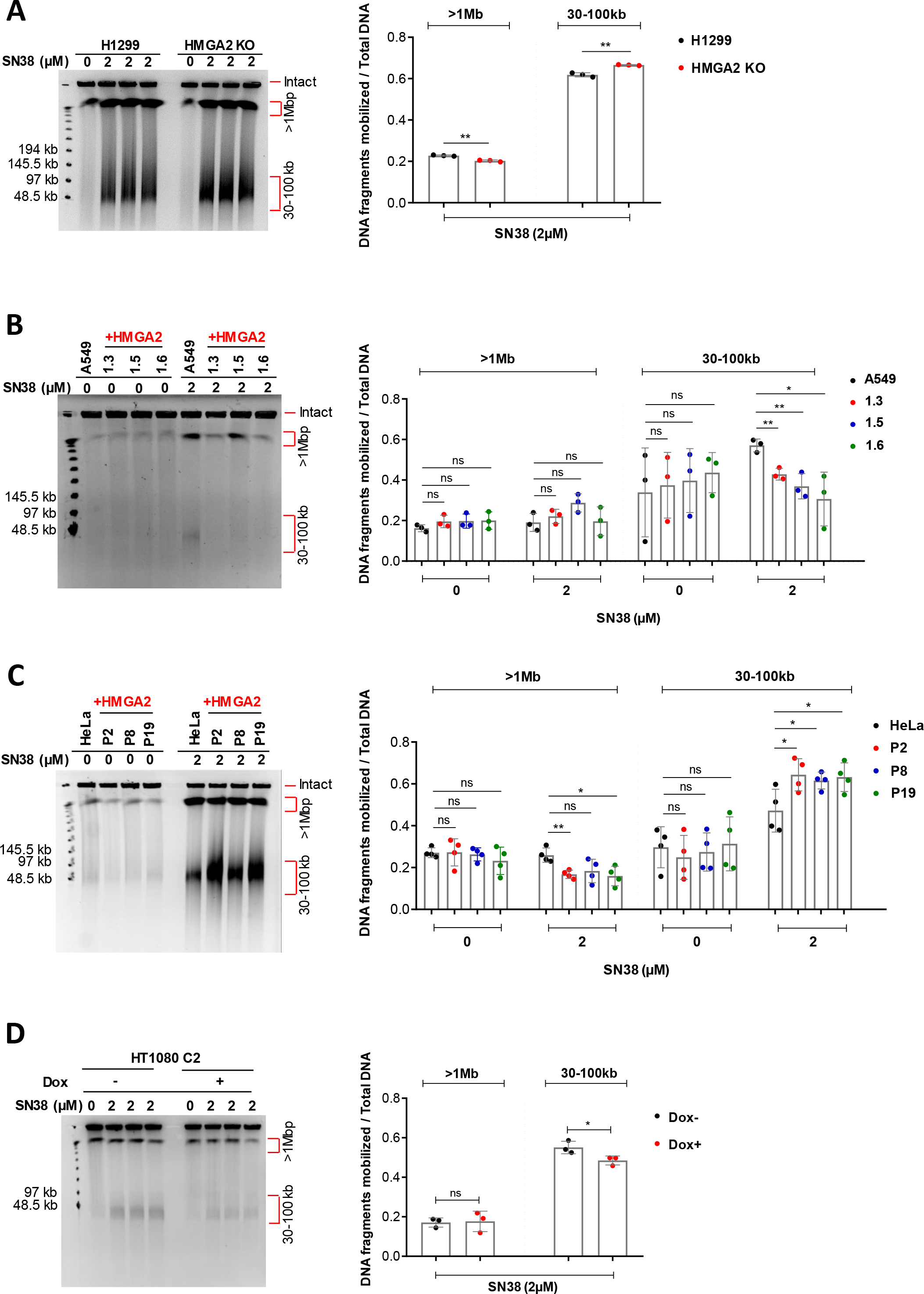
HMGA2 expression controls sensitivity to TOP1 poison SN38. **(A)** PFGE analysis of DSB formation in H1299 cells (parental and HMGA2 KO) in response to 48h incubation with SN38 (left). Quantification of SN38-induced DNA fragments (>1Mb and 30-100kb fractions) was done by ImageJ software (right) with each fragment fraction normalized to total DNA loaded (n=3 independent experiments). Error bars show s.d. Unpaired two-tailed t-tests. * p < 0.05. **(B)** Representative PFGE analysis of DSB formation in A549 cells (parental and three recombinant HMGA2-expressing cell lines) in response to 48h incubation with SN38 (left). Quantification of SN38-induced DNA fragments (>1Mb and 30-100kb fractions) (right) was done by ImageJ software with each fragment fraction normalized to total DNA loaded (n=3 independent experiments). Error bars show s.d. Unpaired two-tailed t-tests. * p < 0.05, ** p < 0.01. **(C)** Representative PFGE analysis of DSB formation in HeLa cells (parental and three recombinant HMGA2-expressing cell lines) in response to 48h incubation with SN38 (left). Quantification of SN38-induced DNA fragments (>1Mb and 30-100kb fractions) (right) was done by ImageJ software with each fragment fraction normalized to total DNA loaded (n=4 independent experiments). Error bars show s.d. Unpaired two-tailed t-tests. * p < 0.05, ** p < 0.01, *** p < 0.001. **(D)** Representative PFGE analysis of DSB formation in HT1080 C2 cell line in response to 48h incubation with SN38 (left panel). HMGA2 expression was down-regulated by doxycycline (Dox)-induced shRNA for 96h. Quantification of SN38-induced DNA fragments (>1Mb and 30-100kb fractions) was done by ImageJ software (right panel) with each fragment fraction normalized to total DNA loaded (n=3 independent experiments). Error bars show s.d. Unpaired two-tailed t-tests. ns not significant, * p < 0.05.

**Fig 3.**
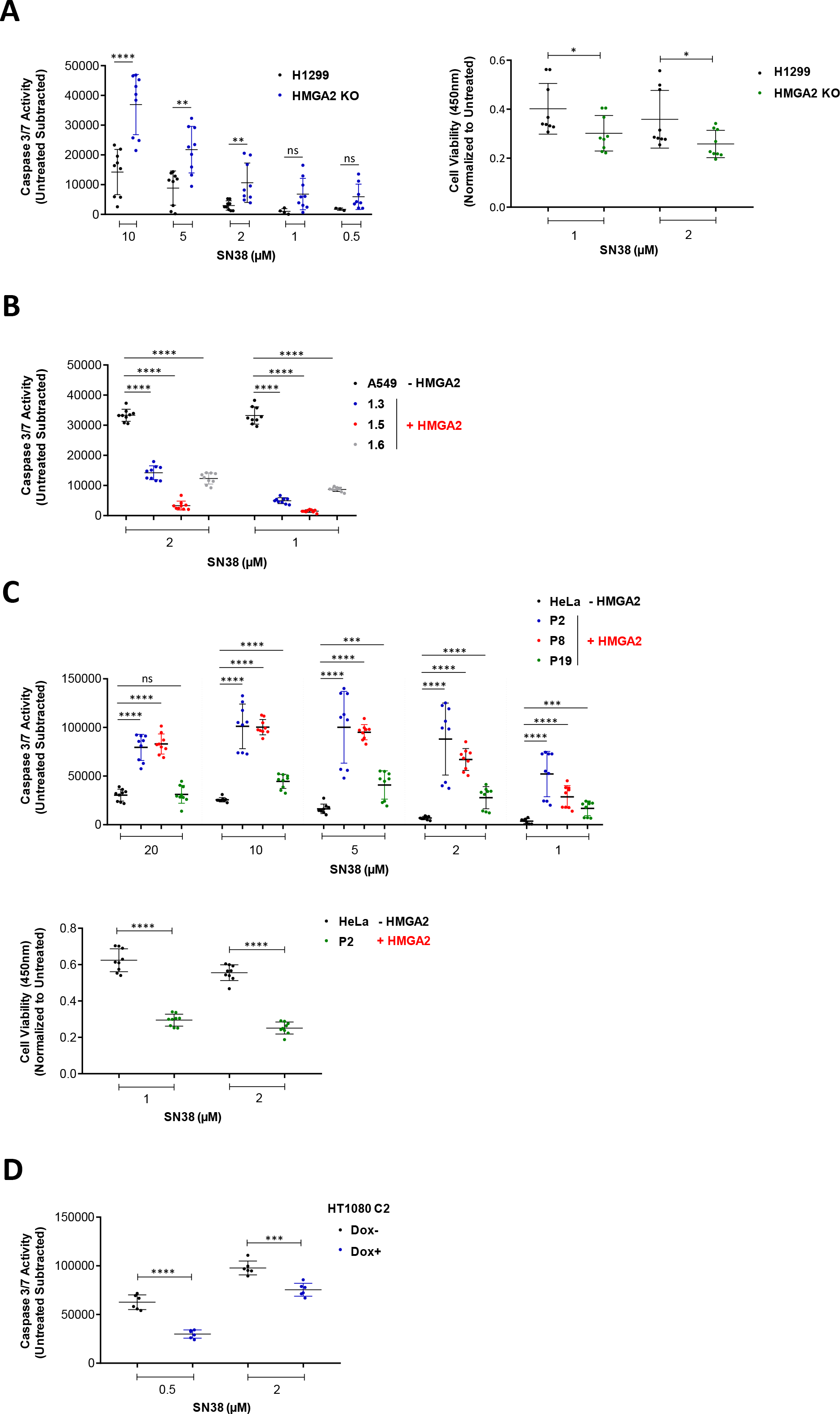
Caspase 3/7 activities correspond with the generation of 30-100kb fragments in response to SN38 treatment. **(A)** Caspase 3/7 activity (left panel) in H1299 cells (parental and HMGA2 KO) after SN38 treatment at indicated doses for 24h and cell survival (CCK8) assay (right panel) with H1299 and HMGA2 KO cells after treatment with SN38 for 48h (n=3 independent experiments with 3 technical replicates in each). Error bars show s.d. Unpaired two-tailed t-tests. * p < 0.05, ** p < 0.01, *** p < 0.001, **** p < 0.0001. **(B)** Caspase 3/7 activity in A549 cells (parental and three recombinant HMGA2-expressing cell lines) after SN38 treatment at indicated doses for 24h (n=3 independent experiments with 3 technical replicates in each). Error bars show s.d. Unpaired two-tailed t-tests. * p < 0.05, ** p < 0.01, *** p < 0.001, **** p < 0.0001. **(C)** Caspase 3/7 activity (top panel) in HeLa cells (parental and three recombinant HMGA2-expressing cell lines) after SN38 treatment at indicated doses for 24h and cell survival (CCK8) assay (bottom panel) with HeLa and P2 cells (recombinant HMGA2 expressing cell line) after treatment with SN38 for 48h (n=3 independent experiments with 3 technical replicates in each). Error bars show s.d. Unpaired two-tailed t-tests. * p < 0.05, ** p < 0.01, *** p < 0.001, **** p < 0.0001. **(D)** Caspase 3/7 activity in HT1080 C2 cells (Dox-/+) after SN38 treatment at indicated doses for 24h (n=2 independent experiments with 3 technical replicates in each). Error bars show s.d. Unpaired two-tailed t-tests. * p < 0.05, ** p < 0.01, *** p < 0.001, **** p < 0.0001.

We next determined whether the observed chemosensitivity to SN38 across the four cancer cell models could be due to differences in expression levels of the SN38 target TOP1. We found that cellular TOP1 levels were very similar irrespective of HMGA2 expression (**S3 Fig**). In addition, a direct comparison across the four models also revealed very similar TOP1 expression levels (**S3 Fig**).

### Cellular chemosensitivity to SN38 depends on HMGA2 levels

With TOP1 levels highly comparable across the tested cell lines, we next asked whether HMGA2 expression levels could be a determining factor in the observed chemosensitivity. Western blot analysis revealed that SN38 resistant H1299 cells (tri-allelic HMGA2 expression) and A549 1.3 cells exhibited significantly higher HMGA2 expression levels compared to drug sensitive HT1080 C1/C2 and HeLa P2 cells (**Fig 4A**). This allowed us to propose a functionally important threshold level, which potentially distinguishes high from low/moderate HMGA2 levels (dotted line in **Fig 4A**). Interestingly, the two cell lines more resistant to SN38 also exhibited higher expression levels of the related HMGA1 protein when compared to the very similar expression levels across the other cell lines (**S3 Fig**). There is evidence that HMGA1 also has an intrinsic replication fork protection function which, could be a contributing factor to the chemosensitivity of cancer cells (27). Taken together, these data raised the intriguing possibility that the amount of HMGA2, which is exclusively nuclear (27), could be a determining factor for SN38 sensitivity.

**Fig 4.**
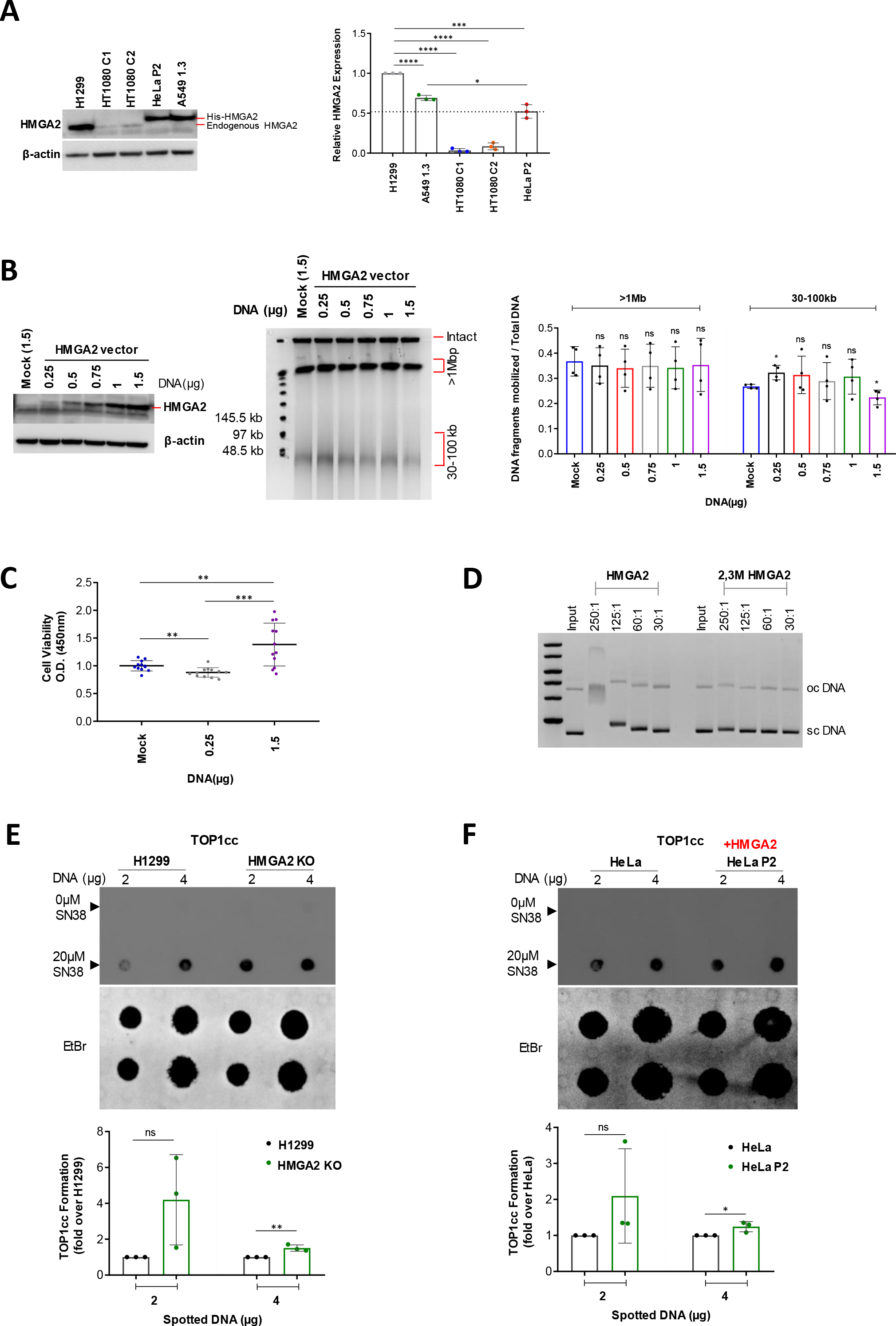
HMGA2 expression levels determine differential chemosensitivity to SN38. **(A)** Comparative Western blot analysis of endogenous (H1299 and HT1080C1/C2 cells) and His-tagged (HeLa P2 and A549 1.3 cells) exogenous HMGA2 levels across various cell lines (left panel). β-actin was used as a loading control. Quantification (right panel) of HMGA2 expression relative to H1299 using ImageJ software (n=3 independent experiments). The dotted line indicates a functional threshold level for high versus low/moderate HMGA2 expression. Error bars show s.d. Unpaired two-tailed t-tests. * p < 0.05, ** p < 0.01, *** p < 0.001, **** p < 0.0001. **(B)** Western blot of HMGA2 expression (left panel) after transfection with indicated amounts of HMGA2 expression vector in H1299 HMGA2 KO cells (mock vector served as control). Representative PFGE analysis of DSB formation in transfected cells after SN38 treatment (2μM) for 48h (centre panel). Quantification of SN38-induced DNA fragments (>1Mb and 30-100kb fractions) (right panel) was done by ImageJ software with ratios (each fragment fraction per intact DNA) plotted as fold change relative to mock transfected cells (100%) for each experiment (n=4 independent experiments). Error bars show s.d. Unpaired two-tailed t-tests. * p < 0.05, ** p < 0.01, *** p < 0.001, **** p < 0.0001. **(C)** Cell survival (CCK8) assay in H1299 HMGA2 KO cells transfected with 0.25μg (low) and 1.5μg (high) amounts of HMGA2 expression vector (mock vector served as control) after SN38 treatment (2μM) for 48h. Data plotted as fold change with mock as 100% (n=4 independent experiments with 3 technical replicates in each). Error bars show s.d. Unpaired two-tailed t-tests. * p < 0.05, ** p < 0.01, *** p < 0.001. **(D)** Representative complex formation of HMGA2 and 2,3M HMGA2 carrying mutated AT-hooks 2 and 3 with supercoiled plasmid DNA at indicated HMGA2:DNA molar ratios. HMGA2-DNA complexes were subjected to electrophoresis in 0.8% agarose gels and DNA was stained with ethidium bromide (n=2 independent experiments). Positions of sc (supercoiled) and oc (open circular/nicked) DNA are indicated. **(E)** Representative *In vivo* complex of enzyme (ICE) assay (see **Methods**), comparing the intracellular formation of TOP1cc between H1299 and HMGA2 KO cells after 20μM SN38 treatment for 30mins. The amount of genomic DNA (μg) loaded is indicated (top) and DMSO-treated cells were used as controls. Quantification of SN38 induced TOP1cc formation (right) with intensities normalized to amounts of spotted DNA that were visualized by staining with Ethidium bromide and plotted as fold change relative to parental H1299 cells (n=3 independent experiments). Error bars show s.d. Unpaired two-tailed t-tests. ns not significant, * p < 0.05, ** p < 0.01. **(F)** Representative *In vivo* complex of enzyme (ICE) assay, comparing the intracellular formation of TOP1cc between HeLa and HeLa P2 cells after 20μM SN38 treatment for 30mins. DMSO treated cells were used as controls. Quantification of SN38 induced TOP1cc formation (right) with intensities normalized to the amounts of spotted DNA that were visualized by staining with Ethidium bromide and plotted as fold change relative to parental HeLa cells (n=3 independent experiments). Error bars show s.d. Unpaired two-tailed t-tests. ns not significant, * p < 0.05.

To rule out differential HMGA2 expression due to drug effects, we treated the high HMGA2 expressing H1299 cells with SN38 for 48h and found no alterations in HMGA2 expression levels (**S3 Fig**). We next tested directly whether HMGA2 expression levels determine SN38 chemosensitivity and performed complementation assays by transfecting various amounts of HMGA2 expression vector into H1299 HMGA2 KO cells (**Fig 4B**). We found that the generated amounts of 30-100kb fragments increased at low but decreased upon very high HMGA2 expression. As expected, cell viability assays revealed that low HMGA2 levels led to reduced cell survival, whereas high HMGA2 levels resulted in increased cell survival compared to mock control **(Fig 4C)** These data confirm that the SN38 treatment outcome depended on HMGA2 expression levels. Additionally, we asked whether the high expression levels of HMGA2 in H1299 cells could lead to differences in cell proliferation, which, in turn, could affect SN38 chemosensitivity. We found that in comparison to H1299 wild-type cells which express high levels of HMGA2, KO cells did not show significant differences in cell growth (**S3 Fig**). Taken together, the observed differential chemosensitivity against SN38 suggests that HMGA2’s fork chaperone function (27) is, at least in part, mechanistically connected to the formation of supercoiled DNA during replication which is a TOP1 substrate.

### Chemosensitivity to SN38 correlates with HMGA2-mediated TOP1cc formation

Our recent *in vitro* studies had shown that HMGA2 efficiently scrunches (+/−) plectonemic supercoils through high affinity DNA segment bridging, leading to extended, rod-like HMGA2-scDNA structures (28). We employed gel retardation assays with purified wild-type and variant HMGA2 to confirm that AT-hooks were necessary for compact and stable higher-order HMGA2-scDNA complex formation (**Fig 4D**). Furthermore, our recent biochemical assays had shown that at a moderate range of HMGA2:scDNA stoichiometries, SN38-induced TOP1cc formation was enhanced 2- to 3-fold. At high stoichiometries, however, HMGA2 negatively interfered with TOP1 supercoil relaxation and reduced TOP1cc formation (29). It is likely that this results from direct competition between HMGA2 and TOP1, as both proteins recognize plectonemic DNA segment crossings in scDNA (28, 44). These *in vitro* data raised the intriguing possibility that, depending on HMGA2 levels, the protein is able to affect SN38 binding that can lead to either more or less productive drug-TOP1 interactions in ternary complexes (29). In line with these biochemical studies, we noted that the SN38-sensitive HeLa-P2 and HT1080-C1/C2 cells exhibited significantly reduced HMGA2 levels compared to the SN38-resistant H1299 and A549-1.3 cells (**Fig 4A**). Using monoclonal antibodies raised against TOP1cc (45), dot-blots with genomic DNA isolated from cells after a short (30 minutes) exposure to SN38 revealed that the high HMGA2 levels in H1299 cells correlated with reduced amounts of TOP1cc when compared to H1299 HMGA2 KO cells. In contrast, the comparatively moderate HMGA2 levels seen in HeLa-P2 cells correlated with an accumulation of TOP1cc in comparison to HeLa cells which lack detectable HMGA2 protein expression (**Fig 4E and 4F**).

### HMGA2 potentiates SN38-induced inhibition of TOP1 on (+) scDNA

A previous single DNA nanomanipulation study showed that the camptothecin drug topotecan significantly hindered TOP1-mediated uncoiling, and, surprisingly, that this inhibitory effect was particularly pronounced on (+) supercoiled DNA (18). Using a similar approach with magnetic tweezers and torsionally constrained single DNA molecules (the experimental set-up is schematically introduced in **S4 Fig**), we investigated the effect of human TOP1 on both geometric forms of supercoiling. We confirmed that strong inhibition of relaxation of (+) supercoiling is also observed with the irinotecan derivative SN38 (**Fig 5A and 5B**; controls without SN38 and HMGA2 are shown in **S4 Fig**). Interestingly, while the presence of 500 nM HMGA2 only moderately impaired the relaxation activity of human TOP1 on both (−) and (+) supercoiled DNA (**Fig 5C and 5D**), the combined effects of HMGA2 and SN38 on relaxation were much greater than additive, and were particularly pronounced on (+) supercoiled DNA (**Fig 5E, 5F and 5G; also see S4 Fig for a zoomed in graph**).

**Fig 5.**
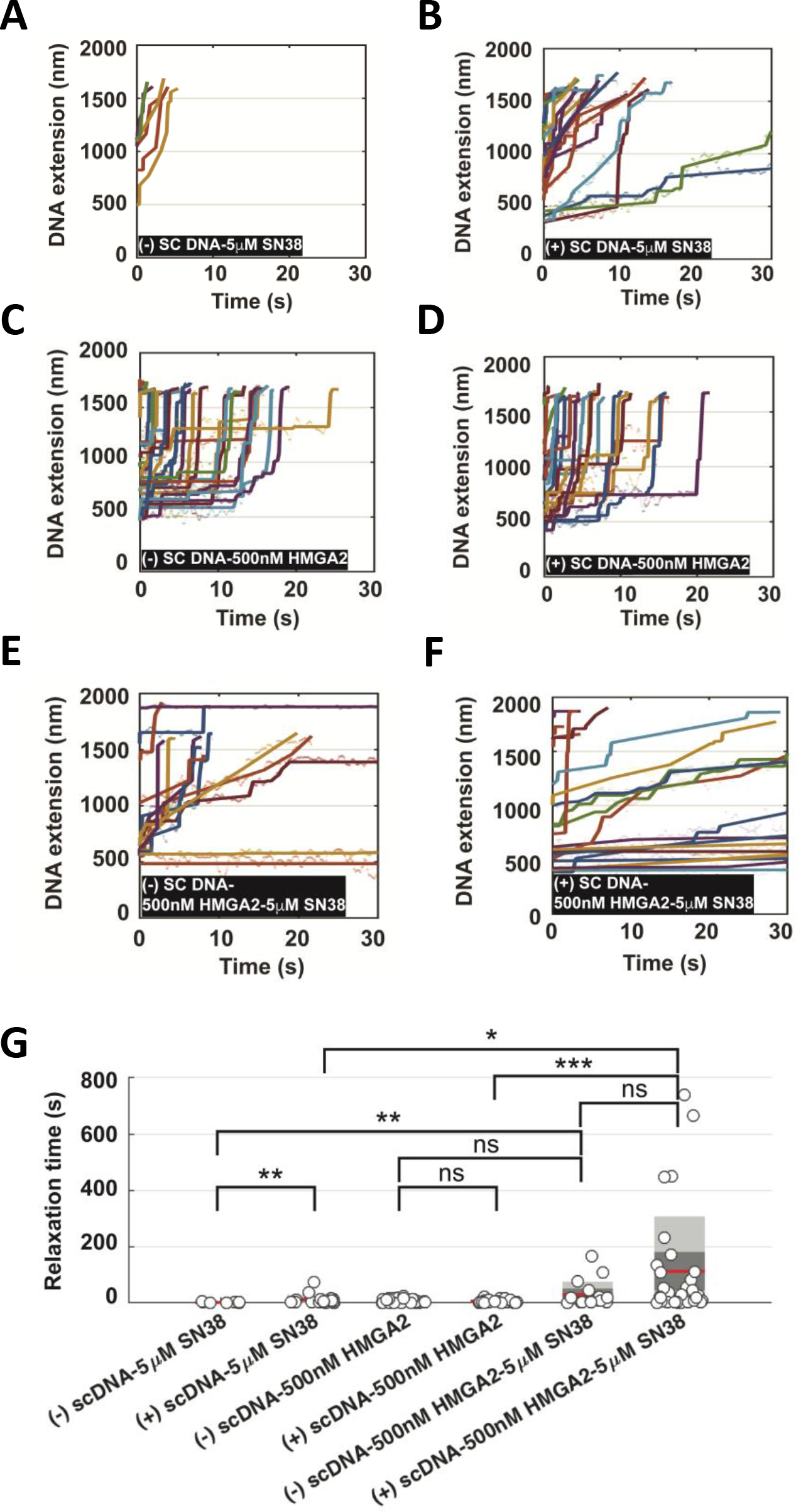
Synergistic effects of SN38 and HMGA2 on supercoil relaxation by human Topoisomerase I. **(A-B)** In the presence of 5nM topoisomerase 1 and 5μM SN38, fast relaxation of (−) scDNA and slow relaxation of (+) scDNA are observed indicated by segmented extension increases linearly. The scatter plot of the raw data is fitted by piecewise linear regression (also see **S4. Fig**). **(C-D)** In the presence of 5nM topoisomerase 1 and 500nM HMGA2, both (−) scDNA and (+) scDNA are relaxed in a jump-pause manner for a similar duration. **(E-F)** In the presence of 5nM topoisomerase 1, 5μM SN38 and 500nM HMGA2, many of the relaxation events are greatly impeded. The mixed features of extension increase (linear and pause-jump) can be observed. **(G)** Box plot of relaxation time (grey circle) summarized from **(A-F)**. The mean is shown by the red line, grey and dark grey areas represent the distribution of s.d and s.e.m. p-value is calculated by Wilcoxon rank-sum test. A different scale of the same plot is shown in **S4. Fig.** in order to highlight the different relaxation times due to the sole presence of either SN38 or HMGA2.

These single DNA nanomanipulation data in combination with our previously reported results obtained with purified components in biochemical assays (29) provide a mechanistic model proposed in the Discussion section for our current findings with cell-based assays, i.e. that moderate HMGA2 levels could lead to more productive drug-TOP1 interactions in TOP1-(+)scDNA-HMGA2 ternary complexes. Collectively, our data thus argue that the observed differential chemosensitivity of cancer cells to SN38 is caused, at least in part, by the extent of intracellular TOP1cc formation, which is modulated by HMGA2 within ternary scDNA complexes.

### Subtelomeric regions are highly sensitive to topological challenges mitigated by HMGA2

Our results indicated that HMGA2 is critically involved in the control of the generation of distinct 30-100kb genomic DNA fragments during replication stress. This led us to determine the genomic origins of these DNA segments. Corresponding fragments from both SN38-treated high HMGA2 expressing H1299 wild-type and H1299 HMGA2 KO cells (also see **Fig 2A**) were gel extracted and separately subjected to deep sequencing; total genomic DNA from corresponding untreated cells served as controls (**S5 Fig**). Strikingly, similar subtelomeric regions in human chromosomes were significantly enriched in the two independently SN38-treated samples (**Fig 6**; **also see S5 Fig**). We did not detect obvious correlations between the corresponding DNA fragment breakpoint regions, their GC content or the orientation of known nearby gene transcription units.

**Fig 6.**
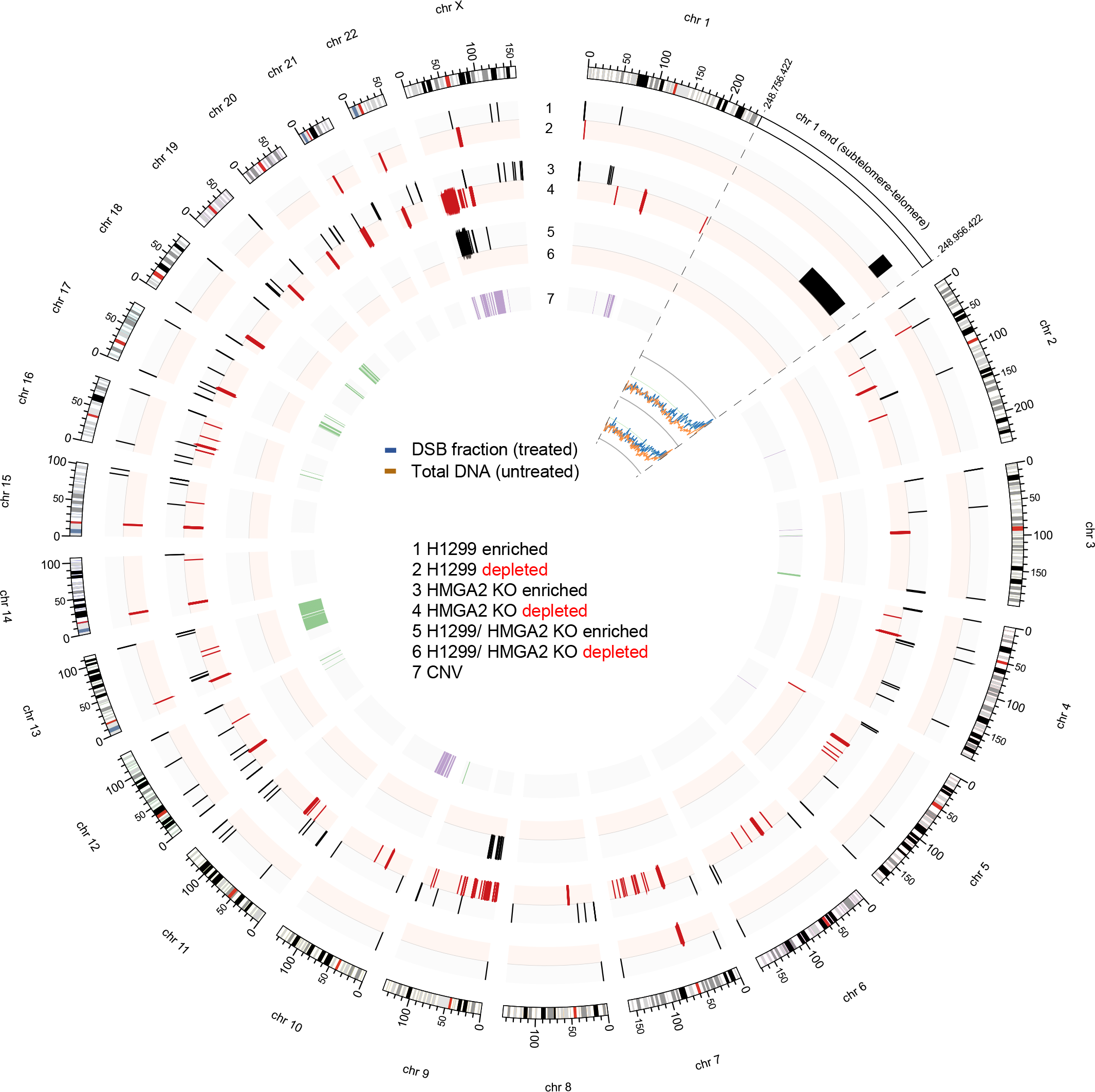
Genome mapping of the SN38 induced double strand break (DSB) fractions. Circos plot showing the genome-wide distribution of differential enrichment and depletion in the SN38-induced 30-100kb fragment fraction of H1299 (plots 1 and 2) and HMGA2 KO (plots 3 and 4) cell lines determined by deep sequencing and normalized to total genomic DNA from untreated cells (plots 1-6 using 20kb bin size and a false discovery rate (FDR) <1%). Subtelomeric regions of the majority of chromosomes were enriched in the 30-100 kb fraction of both SN38-treated cell samples. As an example, we enlarged (500bp bin size) the same region of chr1 for both samples. Similar enrichment of subtelomeric sequences in the 30-100 kb fraction (in blue) when compared to the untreated control DNA (in orange) is apparent for both samples. Copy number variations (CNV) in untreated HMGA2 KO cells compared to untreated parental H1299 cells are shown on plot 7, with purple indicating gain and green indicating loss. The statistical significances of enrichments and depletions are indicated in the center of the plot.

In order to corroborate these results, we performed Southern Blots (SBs) to detect telomeric repeat regions using a specific probe and showed that the high-level endogenous expression of HMGA2 in H1299 parental cells as well as complementation of H1299 HMGA2 KO cells with high exogenous HMGA2 expression levels substantially protected subtelomeres from SN38-induced DSBs (**Fig 7A and 7B**). Likewise, Southern blots performed with DNA extracted from parental and the three HMGA2-expressing HeLa cell lines after SN38 treatment (the corresponding EtBr stained PFGE is shown in **Fig 2C**) confirmed that moderate HMGA2 expression levels increased DSB formation in subtelomeric regions (**Fig 7C**). Intriguingly, the SBs obtained with unchallenged HeLa cells also showed that HMGA2 had a stabilizing effect on subtelomeres (**Fig 7C**; **untreated (30-100kb fractions)**). We conclude that these genomic regions are already fragile, possibly due to endogenous replication stress akin to that induced exogenously by HU (compare **Fig 1A**), and that HMGA2 can mitigate these endogenous challenges. We also found that subtelomeres in human embryonic stem cells (hESCs), which express HMGA2 at low levels comparable to HT1080 cells, are extremely vulnerable to SN38 (**Fig 7D**). By the same approach, we provide evidence that the subtelomeres in the 30-100kb genomic fragment fractions detected via EtBr staining after HU treatment (the corresponding EtBr stained PFGE is shown in **Fig 1A**) were protected by HMGA2 in HeLa cells (**Fig 7E**). Furthermore, we employed the compound TMPyP4, which induces replication fork stalling by stabilizing so-called G-quadruplex DNA secondary structures specifically in G-rich sequences which are present in telomere and subtelomeric regions (46, 47), and found that fractions of 30-100kb fragments enriched in telomere sequences are generated in a dose-dependent manner during 24h TMPyP4 treatment of HeLa cells (**Fig 7F**).

**Fig 7.**
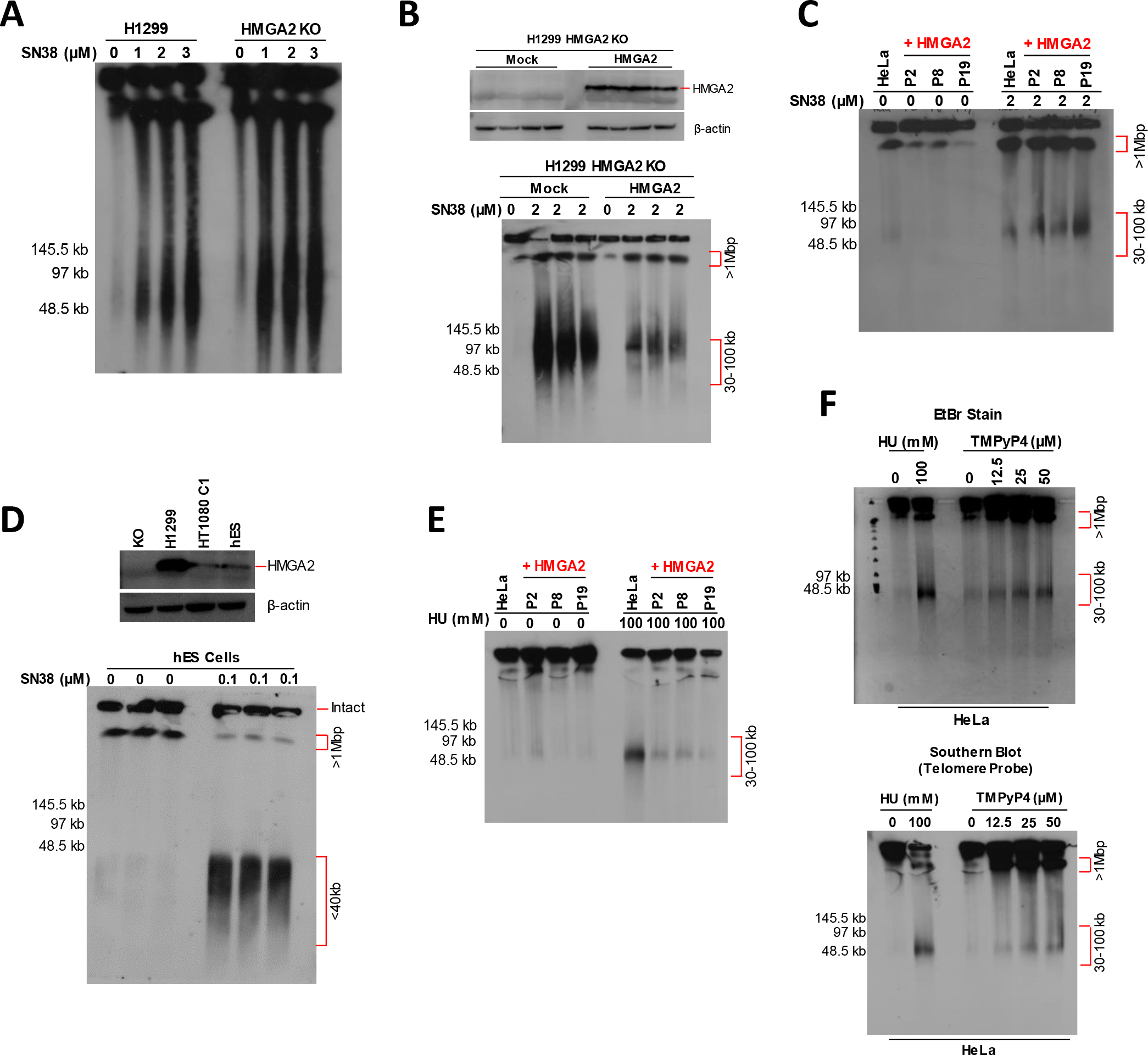
Expression of HMGA2 affects human subtelomeric domains upon induced replication stress. **(A)** Southern blot analysis of H1299 cells (parental and HMGA2 KO cell line) after PFGE separation using telomere-specific probes. SN38 treatment at three different doses was done for 48h, with DMSO used as control. **(B)** Western blot showing exogenous expression of HMGA2 (top panel) after transfection with 1.5μg HMGA2 expression vector in H1299 HMGA2 KO cells (mock vector served as control). Southern blot analysis of HMGA2 KO transfected cells after PFGE separation using telomere-specific probes (bottom panel). 2μM SN38 treatment was done for 48h, with DMSO used as control (n=3 independent experiments). **(C)** Representative southern blot analysis of HeLa cells (parental and three recombinant HMGA2-expressing cell lines) after PFGE separation using telomere-specific probe. 2μM SN38 treatment was for 48h, with DMSO as control (n=3 independent experiments). **(D)** Western blot showing HMGA2 expression for indicated cell lines with β-actin used as internal control (top panel). Southern blot analysis of human embryonic stem (hES) cells after PFGE separation using telomere-specific probes. 0.1μM SN38 treatment was for 24h, with DMSO as control (n=3 independent experiments). **(E)** Representative southern blot analysis of HeLa cells (parental and three recombinant HMGA2-expressing cell lines) after PFGE separation using telomere-specific probe. 100mM HU treatment was done for 24h (n=3 independent experiments). **(F)** PFGE analysis of DSB formation in HeLa cells in response to 24h incubation with TMPyP4 (top panel) and Southern blot analysis after PFGE separation using telomere-specific probes (bottom panel). 100mM HU treatment for 24h was used as control for comparing telomeric DNA fragments (>1Mb and 30-100kb fractions).

### HMGA2 affects chemosensitivity to TOP1 poisons *in vivo*

Irinotecan is widely used in clinical practice to treat colon cancer patients (48), and we next asked whether expression of HMGA2 affected chemosensitivity to irinotecan in patient-derived xenograft (PDX) models of colorectal cancers. SCID mice were inoculated subcutaneously with passaged tissues derived from primary tumors of consented patients. Five PDX models (labeled 935, 1017, 1030, 1073 and 1414) were used, and mice were randomized based on tumor volumes and body weights into vehicle and irinotecan groups (n=6/group). Both groups received corresponding treatments for six consecutive weeks.

PDX models 935, 1073 and 1030 expressed HMGA2 at levels comparable to or lower than those in SN38-sensitive HT1080-C1 cells; models 1017 and 1414 lacked detectable HMGA2 protein expression (**Fig 8A**; **also see S6 Fig**). Strikingly, an overall regression of tumor volumes over a period of 39 days was only observed in HMGA2-positive PDX models (**Fig 8B**). There were no significant differences between the tumor volumes of vehicle groups, indicating that expression of HMGA2 did not affect physiological tumor growth *per se* (**Fig 8C**). The latter result is in agreement with our cell proliferation data comparing H1299 with H1299 HMGA2 KO cells (**S3 Fig**). Taking into account the growth kinetics of each of the models’ own vehicle group, we analyzed the tumor growth inhibition (TGI) percentage at day 39. A TGI of 100% or more indicates tumor regression, and our analysis confirmed that overall regression was only observed in HMGA2-positive PDX models (**Fig 8D and 8E**). These *in vivo* data indicated that low to moderate expression levels of HMGA2 sensitized the PDX models to irinotecan, thus corroborating our *ex vivo* data. High protective HMGA2 levels, which would be comparable to those in SN38-resistant H1299 cells, were not detected in our PDX models (**Fig 8A**).

**Fig 8.**
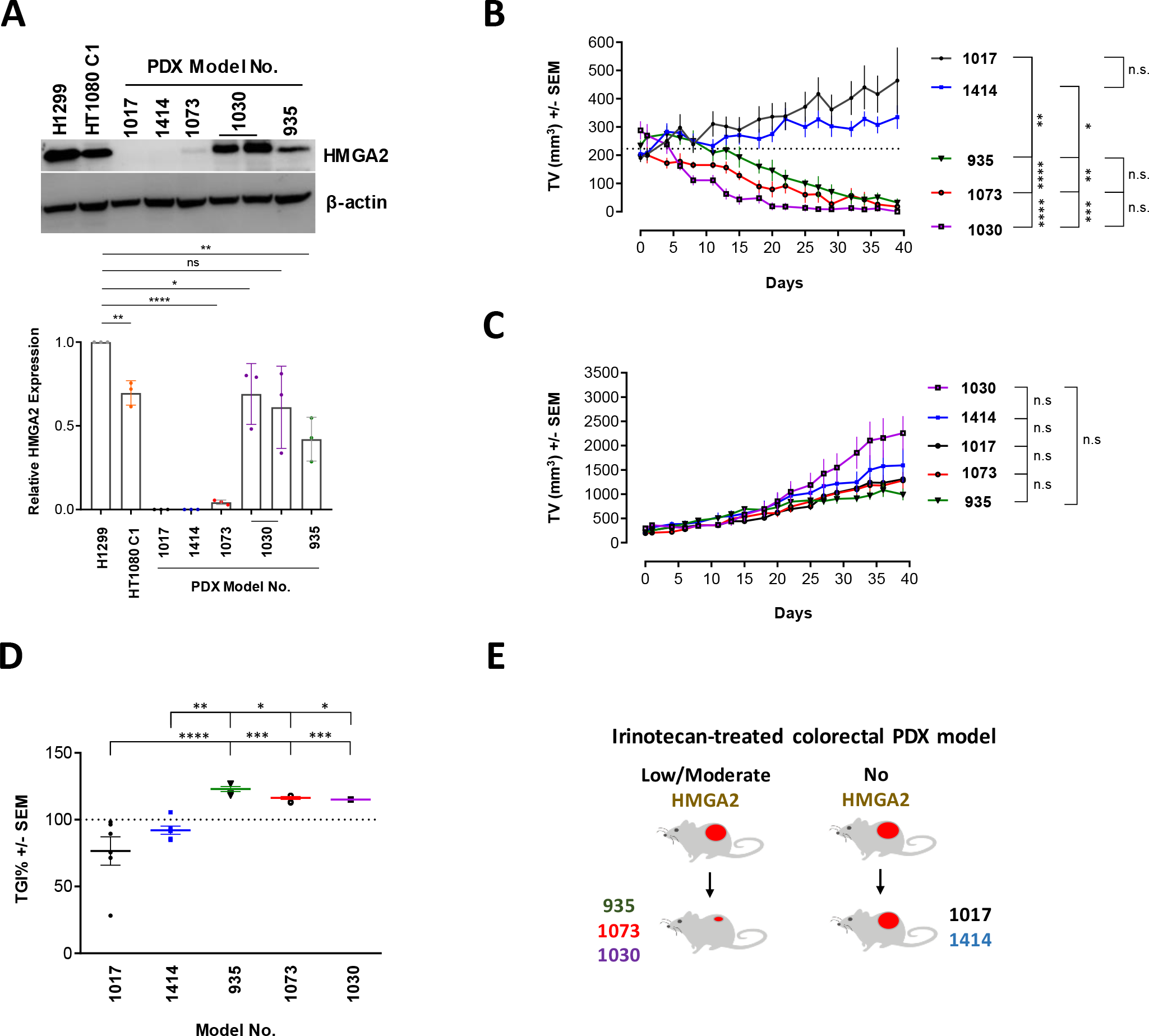
Low-to-moderate HMGA2 expression increases irinotecan chemosensitivity of PDX models. **(A)** Western blot showing HMGA2 expression from representative tumors from each irinotecan-treated PDX model (two samples are from model 1030). We compared these expression levels with those in two HMGA2-expressing cancer cell lines that can serve as standards for SN38 sensitivity: H1299 cells, which exhibit a high degree of SN38 resistance due to very high HMGA2 levels, and HT1080 C1 cells, which exhibit an increased SN38 sensitivity due to moderate HMGA2 levels (see also **S6. Fig.**). Quantification (right) of relative HMGA2 expression using ImageJ software (n=3 independent experiments). Error bars show s.d. Unpaired two-tailed t-tests. * p < 0.05, ** p < 0.01, **** p < 0.0001. **(B)** Tumor growth kinetics (Day 0 to Day 39) of five PDX models (n=6 animals/model) in Irinotecan treatment group. Dotted lines indicate mean starting volume of all PDX models. Two-way ANOVA followed by Tukey’s multiple comparisons. Error bars show s.e.m. * p < 0.05, ** p < 0.01, *** p < 0.001, **** p < 0.0001. **(C)** Tumor growth kinetics (Day 0 to Day 39) of five PDX models (n=6 animals/model) in vehicle group. Two-way ANOVA followed by Tukey's multiple comparisons. Error bars show s.e.m. ns not significant. **(D)** Tumor growth inhibition (TGI) of each PDX model at day 39 (Plotted as percentage). TGI is calculated taking the physiological growth of the tumor (vehicle group) into account. Dotted lines indicate 100% TGI level. Two-way ANOVA followed by Tukey's multiple comparisons. Error bars show s.e.m. * p < 0.05, ** p < 0.01, *** p < 0.001, **** p < 0.0001. **(E)** Graphical representation of tumor outcomes due to irinotecan treatment in colorectal PDX models used in the study.

## Discussion

The effectiveness of cancer chemotherapy is influenced by several factors, with drug resistance and tumor recurrence as major obstacles to successful therapy. However, the underlying mechanisms that alter this efficacy are not completely understood and vary between patients and tumor types. Our integrated data set uncovers a novel mechanism that implicates HMGA2 as an independent, critical determinant of drug response.

We found that the chemosensitivity of cancer cells to the clinically important TOP1 poison irinotecan/SN38 can be determined by the expression level of the oncofetal replication fork chaperone HMGA2. While low to moderate levels sensitized cancer cells and potentiated the formation of genotoxic TOP1cc, high HMGA2 expression levels protected cells against SN38, at least in part by mitigating TOP1cc formation. Such varied response was not observed in cells treated with HU, with broad protection observed across all cell models. We discuss first the proposed role of HMGA2 in the chemosensitivity to irinotecan/SN38.

In conjunction with our recent *in vitro* data (28, 29), our *ex vivo* study now provides evidence that the opposing effects, which depend on HMGA2 expression levels, are mechanistically linked to HMGA2’s ability to constrain supercoiled DNA into highly compacted structures. At low to moderate HMGA2 levels, supercoil constraining by HMGA2 within ternary complexes consisting of (+) scDNA, HMGA2 and TOP1 profoundly affected DNA relaxation in the presence of SN38 *in vitro* (**Fig 5G**). We think that this may result in stabilized or increased drug binding to TOP1 within such ternary complexes, perhaps triggered by an impediment of the required rotational movements of the single-stranded DNA end during TOP1-mediated supercoil relaxation. Such an impediment caused by HMGA2 binding to scDNA that has been cleaved by TOP1 could lead to more frequent misalignments of the O-(3⍰-phospho-DNA)-tyrosine intermediate with the 5⍰-OH end of the otherwise more freely rotating cleaved DNA strand (15). This would indirectly augment the effect of SN38 on strand re-ligation and TOP1cc formation. Our proposed mechanism is supported by biochemical *in vitro* assays showing an increase in TOP1cc formation on scDNA with increasing amounts of HMGA2 (29). In a cellular context, we therefore propose a model in which the increased trapping of TOP1 in covalent complexes with parental DNA during replication will primarily lead to more frequent replication fork run-off events due to collisions between helicases/replisomes and TOP1cc. If this occurs at sub/telomeres, further nucleolytic degradation of parental DNA at these run-off sites over time will guillotine sub/telomeres from the rest of the genomic DNA leading to the 30-100kb telomere-containing genomic fragments that we observed in PFGE/Southern blots (**Fig 9A**).

At very high HMGA2 levels, the supercoiled DNA/chromatin domains, and in particular the DNA segment crossing regions within these domains, may be occupied by HMGA2, thus severely limiting access of TOP1 to its scDNA/chromatin substrate and thereby indirectly interfering with SN38 genotoxic actions (**Fig 9B**). This proposed protective mode against SN38 is also supported by our previous single molecule and biochemical *in vitro* studies (28, 29). Furthermore, constraining most of the DNA supercoils in compact HMGA2-DNA complexes when a significant fraction of TOP1 molecules may be trapped by SN38 at other genomic regions, can potentially enhance fork stability by limiting supercoil-driven fork perturbations, at least temporarily until TOP2 enzymes can relax supercoiling. Such a role of HMGA2 as an effective modulator of transient higher-order chromatin structures, with a particularly important role at human subtelomeres (**Fig 9A,B**), is further supported by the protein’s known high mobility within chromatin and the fact that it binds with high affinity to (+/−) scDNA over other double-stranded DNA conformers (28, 49). These features will enable HMGA2 to quickly associate with transient waves of DNA supercoiling as they emerge in chromatin during replication.

**Fig 9.**
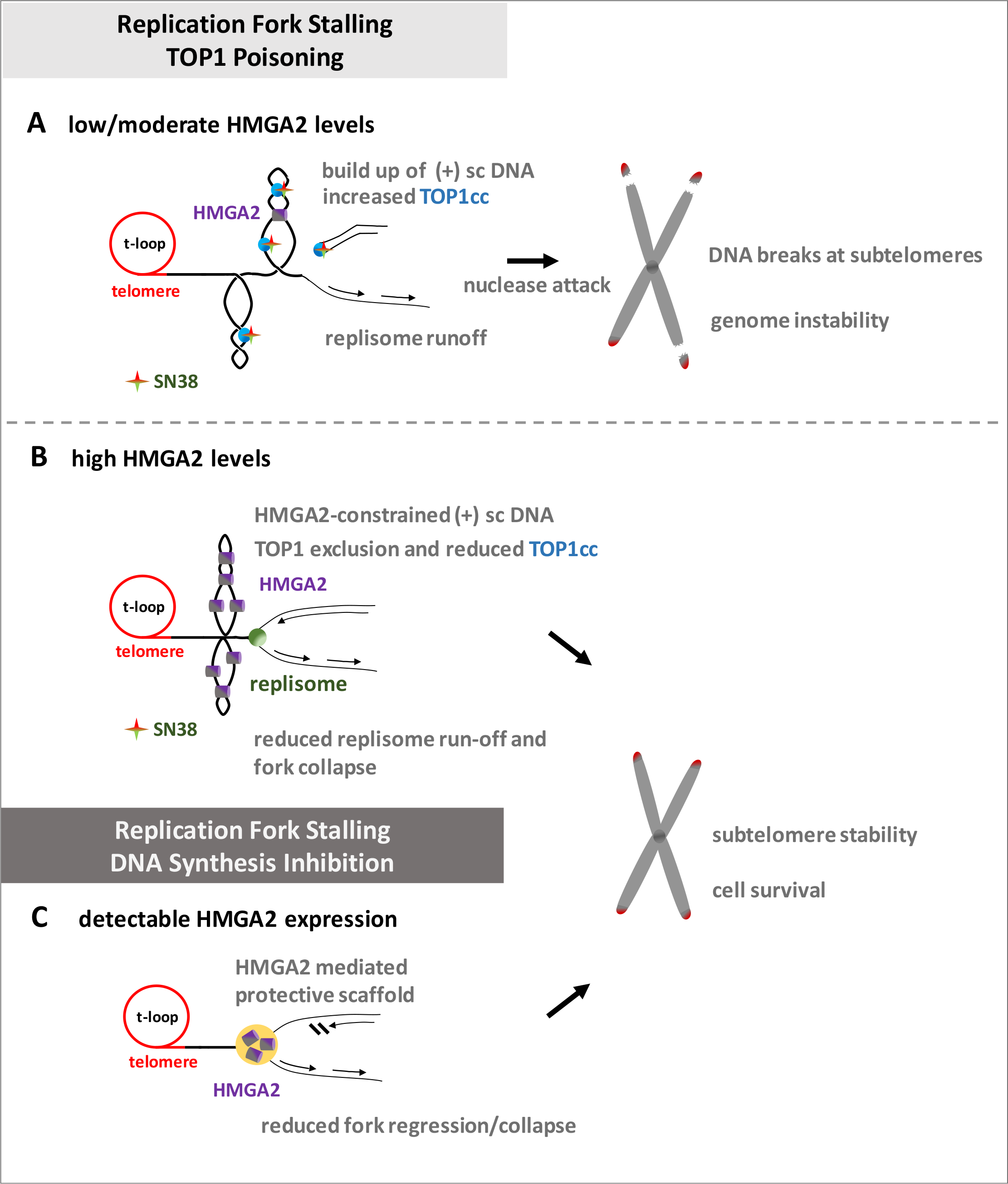
Models for the regulation of subtelomere stability by HMGA2 as a result of replication stress. Replication fork stalling as a result of TOP1 poisoning or DNA synthesis inhibition **(A)** HMGA2 levels determine SN38 treatment responses, with low/moderate HMGA2 levels bound to (+) sc DNA potentiating the accumulation of TOP1-DNA cleavage complexes (TOP1cc), hence leading to more frequent replication run-off events and fork regression/collapse that will ultimately separate sub/telomeres from the rest of chromosomal DNA. **(B)** The accumulation of TOP1cc as a result of SN38 treatment is considerably reduced with high HMGA2 levels, due to constraining of (+) sc DNA and TOP1 exclusion from binding to its scDNA substrate, thereby preventing DSBs at the subtelomeres. **(C)** Such a varied response was not observed with DNA synthesis inhibition by HU since detectable HMGA2 expression always reduced fork collapse and promoted subtelomere stability and cell survival. Described in detail in the Discussion.

An important question is why are the subtelomeric regions identified in our study extraordinarily sensitive to DNA topological challenges induced by SN38? We propose that the proximal canonical [TTAGGG] sequence repeats that make up human telomeres indicate the presence of exceptionally strong DNA topological barriers. It has been argued before that the unique telomeric chromatin composition in human cells, including the deposition of Shelterin complexes and the presence of telomere-loops (t-loops) (**Fig 9A,B**), hinder rapid supercoil dissipation towards the telomeric DNA double strand ends during telomere replication (50–52). It has also been reported that catalytic inhibition of Top2A leads to telomere fragility; a finding that again points at severe DNA topological problems within the telomere repeat regions (53). Furthermore, the Shelterin component Telomere Repeat binding Factor 2 (TRF2) and the exonuclease Apollo work in the same pathway as TOP2A to mitigate the topological consequences of replication stress within telomeres (54). In this context, it is worth noting that replication of most human telomeres originates from within subtelomeric regions (55), hence establishing a unidirectional mode of replication that would prevent rescue of a stalled fork from an incoming second fork that originated within the terminal telomere repeat region. Our data now indicate that strong DNA topological barriers likely exist at the end of human chromosomes and are substantial obstacles to replisomes that translocate along subtelomeric DNA towards the canonical telomere sequences; a process that would inevitably lead to particularly high levels of (+) supercoiling when the supply of TOP1 becomes limiting or when the enzymes are trapped there in TOP1cc. As a consequence, a substantial fraction of forks will run off, and the corresponding DSBs form even at distances up to 100kb or more from the actual telomere repeat sequences (**Fig 9A**). An independent contributing factor to a scenario with impaired supercoil dissipation at the end of our linear chromosomes could be the heterochromatic features of human subtelomeres that were recently identified (56).

The broad protection that HMGA2 provides against HU-induced replication fork collapse into DSBs across all cell systems tested is in line with our previous mechanistic model, which posits that HMGA2 forms a protective DNA scaffold at or nearby stalled replication forks (27) (**Fig. 9C**). Whether temporary constrainment of supercoiling by HMGA2 plays a role in protection against HU-induced fork collapse remains to be investigated further. Because of the possible existence of strong topological barriers at telomeres, as discussed above, such a protective function might be of particular importance to secure genome stability in fast replicating cells. In support of such a scenario, our previous study had revealed that HMGA2 promotes chromosomal stability by reducing the occurrence of HU-induced radials, i.e. structures consisting of multiple chromosomes that are fused at their end regions, perhaps by DSB-induced homologous recombination events (27). Another attractive and mutually not exclusive scenario is that HMGA2 could form a protective scaffold over excess unprotected ssDNA particularly at active firing origins during HU-induced replication stress when most single strand-binding RPA molecules are sequestered (57). Additionally, HMGA2 is capable of binding to secondary DNA structures, such as hairpins, that could form within extended regions of single stranded parental DNA that are generated when a replication fork stalls (35).

This could assist RPA in protecting unwound parental single stranded DNA segments from nucleolytic attack and hence fork collapse into DSBs. In fact, our recent single molecule study revealed that HMGA2 can stabilize hairpins that are formed in single stranded DNA regions (28). In this context, it is noteworthy that our data correlates with a recent extensive *in vivo* study that implicates HMGA2 expression as a prognostic factor to poor clinical outcomes in human pancreatic cancer patients treated with DNA synthesis inhibitors (58).

In conclusion, our study uncovered that certain regions within human subtelomeres appear to be exceptionally vulnerable to DNA topological challenges that result from TOP1 inhibition/trapping, and that the extent of subtelomere instability is influenced by the level of HMGA2. Given that HMGA2 is normally expressed mainly during embryonic/fetal development and aberrantly re-expressed in many human malignancies (26, 59), our integrated study identified an important cancer cell-specific marker for personalized therapeutic interventions with TOP1 poisons and DNA synthesis inhibitors.

## Methods

### Cell culture

Human embryonic stem cells (hESCs) ‘GENEA047’ (Female), were cultured in Genea M2 media as previously described (60). Recombinant HT1080-C1 and HT1080-C2 (male, human fibrosarcoma) cells were previously generated (27) and HMGA2 knockdown was achieved by inducing shHMGA2 with 2μg/ml doxycycline hyclate (Sigma) once every day for four days. Human HMGA2-expressing A549 (male, lung epithelial sarcoma) and HeLa (female, cervical epithelial adenocarcinoma) cells were previously generated (37) and HMGA2 expression was authenticated by western blotting. HT1080, A549 and HeLa cells were cultured in DMEM supplemented with 10% FBS (Gibco), 1% L-Glutamine and 100U/ml each of Penicillin-Streptomycin at 37°C under 5% CO_2_. H1299 (human non-small cell lung carcinoma, ATCC CRL-5803) and HMGA2 KO cells were cultured in RPMI (GIBCO) supplemented with 10% FBS (Gibco) and 100U/ml each of Penicillin-Streptomycin. HMGA2 KO cells were generated using CRISPR-Cas9 by lentiviral infection of HMGA2 sgRNA (5’-ggaggcaggatgagcgcacg-3’) designed to target the immediate downstream region of the start codon in six alleles, thereby creating frame shifts at the N-terminus. Frameshifts were verified by PCR/sequencing. All the cell lines were cultured according to ATCC recommendations, and the cells were not passaged more than ten times from thawing to use. All cell lines have been authenticated by short tandem repeat (STR) genotyping (1st BASE Human Cell Line Authentication Service).

### *In vitro* Gel Retardation Assay

Human recombinant HMGA2 protein and AT-hook 2,3 mutants (27, 29) (in 20mM HEPES pH 7.5, 500mM NaCl, 10% (v/v) glycerol and 2mM TCEP) were isolated from BL21 (DE3) Rosetta cells, using standard techniques that include his-tag affinity chromatography and size exclusion chromatography. Indicated amounts of HMGA2 and mutant HMGA2 protein were incubated with 100ng of negatively supercoiled Renilla reporter plasmid (Promega) for 30mins at 37°C in a buffer containing 10mM Tris-HCl, pH 7.9, 50mM KCl, 50mM NaCl, 5mM MgCl_2_, 0.1mM EDTA, 1mM ATP, 15μg/ml BSA. Samples were mixed with gel loading dye without SDS and analyzed on 0.8% agarose gels in 1X TAE buffer in the absence of ethidium bromide.

### Western blotting

Whole cell lysates were prepared by re-suspending cells in ice-cold RIPA buffer (10mM Tris-HCl pH 8.0, 1mM EDTA, 0.5mM EGTA, 1%Triton X-100, 0.1% Sodium deoxycholate, 0.1% SDS and 140mM NaCl) containing Protease Inhibitor cocktail (Roche) and sonicated on ice (5 W, 10x 3s). Protein concentration was determined using Bio-Rad Protein Assay Dye Concentrate. Samples were heated at 95°C for 10 min and separated by electrophoresis, transferred onto 0.2μm polyvinylidene difluoride membranes (Bio-Rad) and blocked in superblock blocking buffer (Thermo Fisher Scientific). Membranes were incubated with primary antibodies (α-HMGA2 (CST 5269; 1:1000, Ab41878; 1:200), α-HMGA1 (CST 7777S; 1:1000), α-Topoisomerase I (Ab109374; 1:2000), α-β-Actin (Sigma A2228; 1:5000)) overnight at 4°C followed by washing in 0.1%Tween/TBS. Membranes were incubated with appropriate secondary antibodies (Polyclonal goat anti-mouse (Dako, P0447) and polyclonal goat anti-rabbit (Dako, P0448)) at room temperature for 1h and washed three times (10mins each) prior to signal detection. EMD Millipore Immobilon Western Chemiluminescent HRP Substrate (ECL) was used for detection.

### Pulsed Field Gel Electrophoresis

Cells were seeded in a 6-well tissue culture plate and treated with either HU (Sigma) for 24h or SN38 (Abcam) for 6h/48h. Media was changed every 24h during SN38 treatment for 48h in both treated and untreated cells (DMSO control). Harvested cells were embedded in 2% low melting agarose (Sigma) plugs using CHEF disposable plugs (Bio-Rad). After solidification, plugs were incubated in lysis buffer (100 mM EDTA, 0.2% (w/v) sodium deoxycholate, 1% (w/v) sodium lauroyl sarcosine and 1 mg/ml proteinase K; Promega) at 50°C for 24h and washed four times in TE buffer (20mM Tris, pH 8, 50mM EDTA) for 1h each. Plugs were run on 1% megabase agarose (Bio-Rad) on a CHEF DR II equipment (Bio-Rad) under conditions of 120 field angle, 5-30s switch time, 4 V/cm and 14 °C for 14h in 1X TAE. Lambda PFG ladder (48.5-1018kb; NEB) was used as a molecular size marker. Subsequently, DNA was stained with ethidium bromide and quantification was performed using ImageJ (see figure legends for details).

### Southern blotting

Following the resolving of DNA fragments by Pulsed Field Gel Electrophoresis (PFGE), telomeric DNA was detected by Southern blotting using TeloTAGGG Telomere Length Assay kit (Roche, 12209136001) as per the manufacturer’s protocol. Briefly, ethidium bromide stained gels were depurinated in 0.25 M HCl for 10 mins at RT followed by denaturation in 0.5M NaOH and 1.5M NaCl twice at RT for 15 mins each. Subsequently, neutralization was done in 0.5M Tris-HCl, 3M NaCl, pH 7.5 twice at RT for 15 mins each. Further, DNA was transferred onto positively charged nylon membrane (Roche) by overnight capillary transfer at RT using 20X SSC transfer buffer (3M NaCl, 0.3M Sodium citrate, pH 7). DNA was fixed on the membrane by UV-crosslinking at 1.2J/cm^2^ for 30s in HL-2000 Hybrilinker (UVP) and washed twice in 2X SSC. The membranes were prehybridized at 42°C for 1 hour in the ProBlot™ hybridization oven (Labnet) and hybridization was carried out overnight at 42°C using 2μl of the telomeric probe. Membranes were then washed twice (5 mins each) with stringent buffer I (2X SSC, 0.1%SDS) at RT followed by two more washes (20 mins each) with stringent buffer II (0.2X SSC, 0.1%SDS) at 50°C. Following stringent washes, membranes were first incubated in freshly prepared 1X blocking solution for 30 mins before incubating in Anti-DIG-AP working solution for 30 mins. Finally, membranes were washed in 1X washing buffer twice (15 mins each), followed by incubation in 1X detection buffer for 5 mins. Next, substrate solution was added, and the membranes were exposed to X-ray film (Carestream) for 10-20mins which were then developed in X-OMAT 2000 Processor (Kodak).

### Complementation Assay

Lipofectamine 2000 (Invitrogen, 11668-019) transfection reagent was used for all plasmid transfections in H1299 HMGA2 KO cells in 6 well plates. pEF1/Myc-A-hmga2-FLAG expression vector was used to express HMGA2 with pEF1/Myc-A vector as mock control. Prior to transfection, RPMI media was replaced with Penicillin-Streptomycin free media and a 1:2.5 ratio of plasmid DNA(μg): Lipofectamine 2000(μl) was used. Transfection reaction mixtures were vortexed thoroughly and incubated at room temperature for 30mins before adding dropwise to cells. Fresh media containing Penicillin-Streptomycin was added after 6h incubation at 37°C. 36h post transfection, cells were treated with 2μM SN38.

### Cell Survival Assay

Cells were seeded as triplicates in a 96-well black/clear bottom plate and allowed to attach overnight. Cell viability was determined using the cell counting kit-8 (Enzo Life Sciences, ALX-850-039-0100) as per the manufacturer’s instructions. 10μl of CCK-8 solution was added directly to each well after 24h with or without SN38 and incubated for 2hrs at 37°C. Absorbance was measured at 450nm using microplate reader (TECAN Infinite M200 Pro).

### Caspase Activity

5000 cells were seeded as triplicates overnight for each condition in a white-walled 96-well plate and treated with indicated doses of SN38 for 24h. Caspase-Glo 3/7 assays (Promega, G8090) were performed as per the manufacturer’s instructions. Briefly following treatment, Caspase 3/7 activity was measured by adding 100μl of room temperature-equilibrated Caspase/Glo^®^ 3/7 reagent to each well and incubated for 1h at room temperature. Luminescence was measured using a plate reading luminometer (TECAN Infinite M200 Pro).

### *In vivo* complex of enzyme (ICE) assay

Endogenous TOP1cc were detected by Human Topoisomerase ICE kit (Topogen, TG1020-0) following the manufacturer’s instructions. Briefly, cells were seeded in 60mm tissue culture plates and allowed to attach overnight. 20μM SN38 treatment was done for 30mins, following which media was removed and rinsed twice with pre-warmed PBS. Cells were lysed, and the resulting genomic DNA was sonicated at 10% amplitude (3s x4) and diluted in 25mM phosphate buffer, pH 6.5. DNA concentration was estimated using Nano-Drop. Equivalent amounts of DNA were spotted on a Nylon membrane using a dot blot apparatus (Cleaver Scientific) and probed with mouse anti-TOP1cc (EMD Millipore, MABE1084). Ethidium bromide staining of spotted DNA served as loading controls. Quantification was performed using ImageJ (see figure legends for details).

### DNA constructs

The ds DNA (6573bp 48% AT) was obtained by PCR from the 48502 bp phage-λ DNA (New England Biolabs (NEB)) with Q5 Hot start polymerase (NEB). (FP: 5’ – ATTACAAAGTTACCTGTCAAACGGT; RP: 5’ – ACGTAAGGCGTTCCTCGATATG). The two DNA handles (510bp) labeled by multiple digoxigenin and biotin are generated by mixing biotin-16-dUTP and digoxigenin-11-dUTP nucleotides (Roche) with dNTP solution mix in the respective PCR reaction. All three DNA pieces were ligated by incubating them with T4 ligase (NEB) overnight at 16°C.

### Flow channel preparation

Flow channel was first made from two #1 glass coverslips and the bottom one was functionalized by 3-Aminopropyl triethoxy silane (APTES). The channel was then flushed by Silane-PEG-NHS (PG2-NSSL-5k, Nanocs) dissolved in DMSO and incubated for 1 hour. It was followed by flushing anti-digoxigenin Fab fragments (Roche) in the channel to create covalent bonds with NHS group on PEG and incubated for another 1 hour. The channel was then incubated with 1% BSA solution in 1× phosphate buffered saline (PBS) buffer (pH 7.4) at 4°C before experiments to avoid non-specific binding of DNA to the surface.

### Magnetic tweezers experiments

In typical magnetic tweezers experiments, DNA molecules were tethered to a functionalized glass coverslip by a superparamagnetic bead (Dynabeads MyOne) of 1μm in diameter. The force was applied to the DNA molecules through the bead under an inhomogeneous external magnetic field generated by a pair of Neodymium magnets and the force was controlled by the height of the magnets manipulated by a translational micromanipulator (MP-285, Sutter Instruments). A rotation stage (DT-50, Physik Instruments) was used to rotate the magnets in order to wind/unwind torsion-constrained DNA tethers. The change of linking number (Lk) causes DNA twist deformation and results in accumulation of twist elastic energy. Above a threshold, the accumulated energy is relaxed through chiral bending into plectonemic supercoiling conformation at forces below 0.5pN (61–63), which is indicated by the DNA extension decreasing linearly as a function of ΔLk.

Rotating the magnetic bead 30 turns results in winding/unwinding DNA and thus positive (+)/negative (−) supercoiled DNA is formed. Adding 5nM recombinant human topoisomerase 1 (PROSPEC) is capable of removing any onset supercoiled DNA if transiently formed showing that extension remained nearly at the level of relaxed DNA conformation. A mixture of 5nM TOP1 and 5μM SN38 was added into the channel and the SN38 dependent effect was investigated by multiple cycles of relaxation on both (−) scDNA and (+) scDNA. Similar experiments were performed in the presence of 5nM TOP1 and 500nM HMGA2 for both (−) scDNA and (+) scDNA and both scDNA were relaxed in a jump-pause manner for a similar duration. The buffer solution used in the experiments contained 100 mM KCl and 20 mM Tris (pH 7.4). All experiments were conducted at 23 ± 1°C.

### Whole genome sequence analysis

H1299 and HMGA2 KO cells were treated with 2μM SN38 for 48h. Cells from both samples were processed for pulsed field gel electrophoresis. The DNA fragment fraction from each lane, ranging from 30kb up to 150kb was excised after ethidium bromide staining and pooled in order to obtain sufficient amounts for deep sequencing (**S5 Fig**). DNA was extracted using the ZymoClean large fragment DNA recovery kit according to the manufacturer’s instructions. Briefly, Buffer ADB was added to each tube containing the excised gel in the ratio of 3:1 and incubated at 55°C until the gel was completely dissolved. The solution was pooled for each sample and transferred to the Zymo-Spin column followed by washing twice with DNA Wash Buffer. The extracted DNA was eluted using DNA Elution Buffer and quantified using Qubit dsDNA HS Assay Kit (Thermo Fisher Scientific). The quality of DNA was checked by agarose gel electrophoresis. Genomic DNA from H1299 and HMGA2 KO cells were extracted using DNeasy Blood & Tissue Kit (Qiagen) and submitted for sequencing as controls. The extracted genomic DNA was sequenced using Illumina HiSeq High Output sequencer with paired-end sequencing. The read size was 101bp and all four samples were read to depths of at least 300 million reads. The sequencing data was subjected to quality control analysis by FASTQC (64). The sequencing adapters were removed using trimmomatic (65) under default settings. Reads were aligned to GRCh38 using bowtie2 (66) in paired-end mode with default parameters. Reads were binned by bamCoverage using the public Galaxy server (67). Since our fragments ranged between 30-100kb in size, we began with 20kb binning to determine the enrichment in the SN38-induced DSB fraction. False discovery rate (FDR) was calculated using the fdrtool R package (68), and regions under 1% FDR were considered enriched or depleted. UCSC genome browser was used to visualize the coverage for individual tracks, and the genome-wide distribution was plotted using Circos (69).

### *In vivo* animal studies

All animal experiments were performed in the Biological Resource Centre, Agency for Science, Technology and Research (A*STAR). Five to six weeks old SCID female mice (C.B-Igh-1^b^/IcrTac-Prkdc^scid^), 18-22 g at the time of tumor inoculation were purchased from Invivos Pte Ltd (Singapore) and housed in individually ventilated cages in Biological Resource Centre, A*STAR. Room lighting was set to a 12-hours light-dark cycle as recommended by the National Advisory Committee for Laboratory Animal Research (NACLAR). Animals were provided with irradiated Altromin 1324 diet and autoclaved water, ad libitum. Patient derived Xenografts were provided by the Genome Institute of Singapore (GIS), A*STAR. These samples were derived from primary tumor tissues collected from consented colorectal cancer patients from Singapore General Hospital and National Cancer Centre Singapore under protocol 2015/2165 approved by the SingHealth Institutional Review Board. Primary tissues were propagated into immunocompromised mice, and later harvested and bio-banked. Each SCID mouse was inoculated subcutaneously with about 100 mg of PDX tumor material. When tumors reached between 150 mm^3^ to 350 mm^3^, mice were randomized into control and treatment groups (n= 6 mice/group, 3 animals/cage, experimental unit: mouse/tumor). Mice from the control groups received only the vehicle, at a dose volume of 0.1 ml per 10 grams of body weight. Animals in the treatment groups were given Irinotecan (CAS#: 136572-09-3, Active Biochem) at a dose of 50 mg/kg. Both vehicle (PEG 400, Sigma and Water for Injection, BBraun; mixed 1:1 v/v) and drug were administered intraperitoneally, once weekly in the afternoon, for 6 consecutive weeks. Tumor measurements were taken with Vernier calipers three times a week. We calculated tumor volume at two perpendicular axes (L, longest axis; W, shortest axis) using the formula (L X W^2^)/2. Tumor growth inhibition was determined using the formula (C_dayB_ – T_dayB_)/(C_dayB_ – C_dayA_) X 100, where C is the control group, T is the treatment group, dayA is the mean tumor volume on the day of first dose and dayB is the tumor volume at a specified time point. No animals were excluded during this study, and study groups were not blinded. We followed the ARRIVE guidelines, and this study was approved by the Biological Resource Centre IACUC (Institutional Animal Care and Use Committee, protocol #171207).

### Ethics Statement

Patient derived Xenografts were provided by the Genome Institute of Singapore (GIS), A*STAR. These samples were derived from primary tumor tissues collected from colorectal cancer patients with written consent from Singapore General Hospital and National Cancer Centre Singapore. The study was approved by the IRB or Ethics Committee named SingHealth Institutional Review Board under the approval no. 2015/2165. We followed the ARRIVE guidelines, and this study was also approved by the Biological Resource Centre IACUC (Institutional Animal Care and Use Committee, protocol #171207).

## Data and Software Availability

Sequencing data is available at Sequence Read Archive (SRA) as SRR6206333, SRR6206334, SRR6206335 and SRR6206336.

Original data deposited at Mendeley Dataset and is available at: https://data.mendeley.com/datasets/f5t55gkrwz/draft?a=8bba927a-f60a-4a48-8ccd-dd559a8e168a

## Acknowledgements

Special thanks go to Costerwell Khyriem for expert advice on genomic DNA sequence analysis.

## Supporting Information

**S1 Fig. HMGA2 protects against hydroxyurea-induced fork collapse.**

**(A)** A western blot shows corresponding HMGA2 levels (top panel) in HT1080 C1/C2 clonal cell lines. HMGA2 expression was down-regulated by doxycycline (Dox)-induced shRNA for 96h. PFGE analysis of DSB formation in response to 24h incubation with HU (bottom panel). Quantification of HU-induced DNA fragments (>1Mb and 30-100kb fractions) was done by ImageJ software with each fragment fraction normalized to total DNA loaded. These experiments are independent reproductions of those presented in Yu, Lim (27).

**S2 Fig. HMGA2 modulates sensitivity to TOP1 poison.**

**(A)** Representative PFGE analysis of DSB formation in HT1080 C1 cell line in response to 48h incubation with SN38 (left panel). HMGA2 expression was down-regulated by doxycycline (Dox)-induced shRNA for 96h. Quantification of SN38-induced DNA fragments (>1Mb and 30-100kb fractions) was done by ImageJ software (right panel) with each fragment fraction normalized to total DNA loaded (n=3 independent experiments). Error bars show s.d. Unpaired two-tailed t-tests. * p < 0.05, ** p < 0.01, *** p < 0.001, **** p < 0.0001.

**(B)** PFGE analysis of DSB formation in HeLa cells (parental and HMGA2 expressing cell line (P2)) in response to 6h incubation with SN38 (left panel). Quantification of SN38-induced DNA fragments (>1Mb and 30-100kb fractions) was done by ImageJ software (right panel) with each fragment fraction normalized to total DNA loaded (n=3 independent experiments). Error bars show s.d. Unpaired two-tailed t-tests. * p < 0.05, ** p < 0.01.

**S3 Fig. HMGA2 expression levels key to observed differential chemosensitivity to SN38**

**(A)** Western blot showing human TOP1 expression across all tested cell lines (H1299 (parental and HMGA2 KO cells), A549 cells (parental and three recombinant HMGA2-expressing cell lines), HeLa cells (parental and three recombinant HMGA2-expressing cell lines) and HT1080 C1/C2 (Dox-/+) cells).

**(B)** Representative Western blot comparing human TOP1 expression across various cell lines. Quantification (bottom) of human TOP1 expression relative to H1299 cells using ImageJ software (n=3 independent experiments). Error bars show s.d. Unpaired two-tailed t-tests. ns not significant.

**(C)** Representative Western blot comparing HMGA1 expression across various cell lines. Quantification (bottom) of HMGA1 expression relative to H1299 cells using ImageJ software (n=2 independent experiments). Error bars show s.d. Unpaired two-tailed t-tests. ns not significant, * p < 0.05.

**(D)** Western blot showing HMGA2 expression after 2μM SN38 treatment for 48h in H1299 cells (3 technical replicates). DMSO treated cells used as experimental control. β-actin was used as a loading control.

**(E)** Cell survival (CCK8) assay in H1299 and HMGA2 KO cells, analyzed for growth differences up to 4 days (n=2 independent experiments with 3 technical replicates for each time point). Data normalized to 24h time point. Error bars show s.d. Paired two-tailed t-tests. ns not significant.

**S4 Fig. Synergistic effects of SN38 and HMGA2 on supercoil relaxation by human Topoisomerase I**

**(A)** In the absence of SN38, a representative time-trace of extension (top panel) of a torsionally constrained DNA held at 0.3pN during clockwise and anti-clockwise rotation of the bead (bottom panel). The extension decrease suggests that (+/−) supercoiled DNA is generated by the rotating magnetic beads. Inset: sketch of torsionally constrained DNA in the (+/−) supercoiled and relaxed conformation.

**(B)** In the presence of 5nM TOP1, DNA extension remains at the level of unconstrained DNA during bead rotations suggesting that topoisomerase I effectively relaxed supercoiled DNA via ssDNA cleavage activity.

**(C)** In the presence of 5nM TOP1 and 5μM SN38, slow relaxation of positive supercoiled DNA is observed. When the DNA extension is relaxed to its original length, another 30 turns is applied to the DNA and thus cycles of DNA extension-relaxation events are recorded.

**(D)** The representative relaxation event from (C) highlighted in red box is fitted by piecewise linear regression.

**(E)** Box plot of relaxation time (grey circle) summarized for the effects of SN38 and HMGA2 on DNA supercoil relaxation by human topoisomerase I.

**S5 Fig. Experimental pipeline for the mapping of SN38-induced genomic fragments**

**(A)** Analysis of DSB formation by PFGE in H1299 cells (parental and HMGA2 KO) in response to 48h incubation with SN38 (left panel). Quantification of SN38-induced DNA fragments (>1Mb and 30-100kb fractions) was done by ImageJ software (right panel) with each fragment fraction normalized to total DNA loaded (n=4 independent experiments). Error bars show s.d. Unpaired two-tailed t-tests. ns not significant, * p < 0.05, ** p < 0.01, *** p < 0.001. Note the increase of 30-100kb fragments (marked in red) in HMGA2 KO cells which were gel extracted, combined and sequenced for each cell type. **(B-D)** Sequencing Workflow.

**(B)** Scheme of representative PFGE image highlighting extracted fragments (red box).

**(C)** Sequencing reads aligned to GRCh38 and coverage of chr17 is shown as an example.

**(D)** Enrichment ratio for each bin was calculated (treated versus untreated samples), and the data for H1299 is shown (left). Values plotted for chr17 with FDR <0.01 as cut-off for enrichment/depletion (right).

**S6 Fig. Low-to-moderate HMGA2 expression increases irinotecan chemosensitivity of PDX models.**

Western blots showing HMGA2 expression of individual irinotecan treated PDX tumors from each model (horizontal numbers) in comparison with HMGA2 expressing cell lines, as indicated. β-actin was used as a loading control. Note the variation in HMGA2 expression in PDX tumor model 1030.

