## Supplementary figures and images for "The chromatin structuring protein HMGA2 influences human subtelomere stability and cancer chemosensitivity"

### S1 Fig

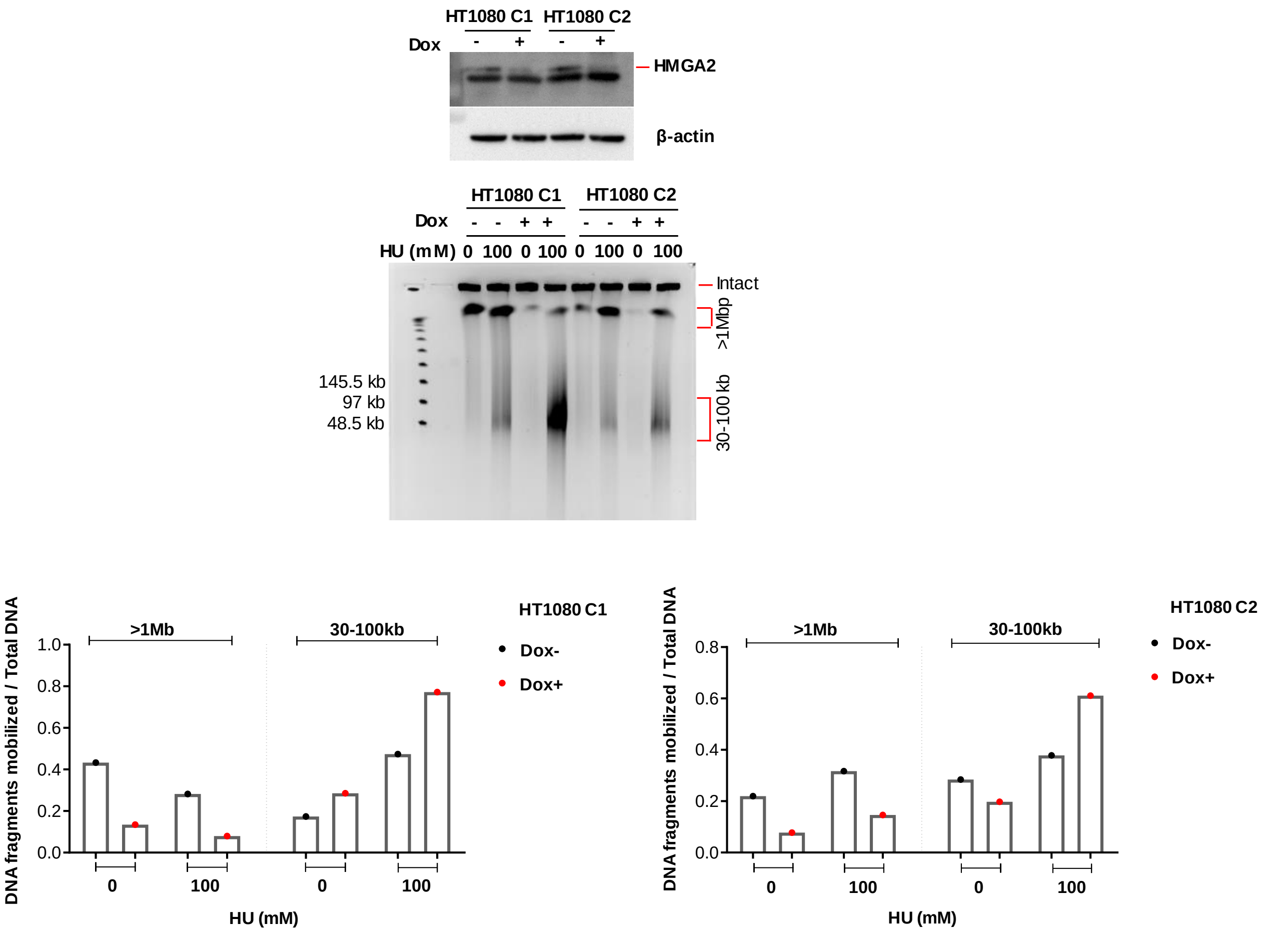

### S2 Fig

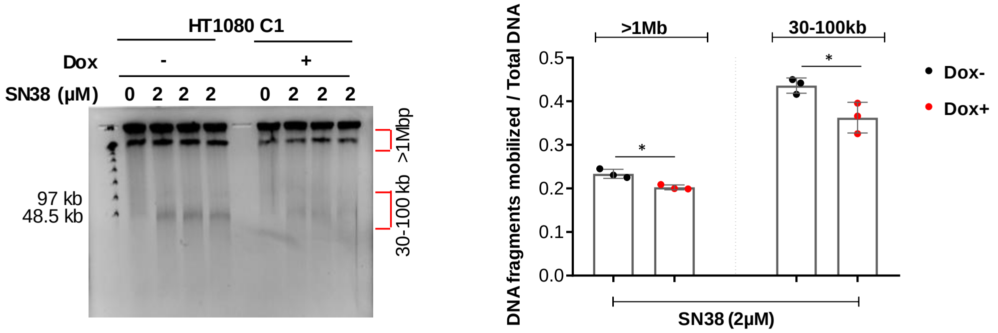

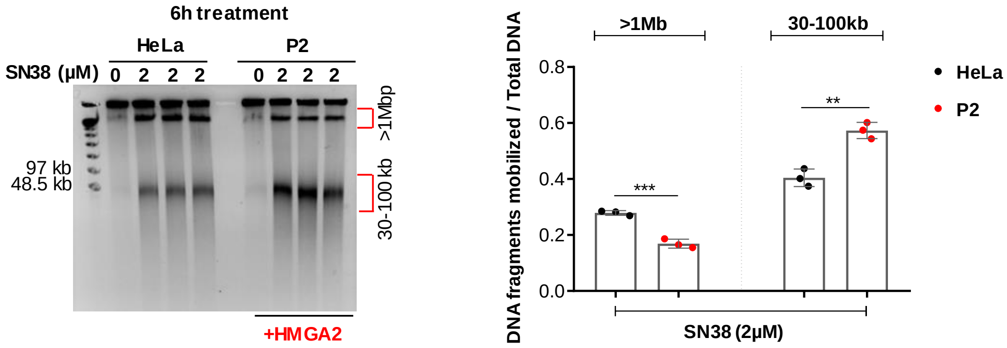

### S3 Fig

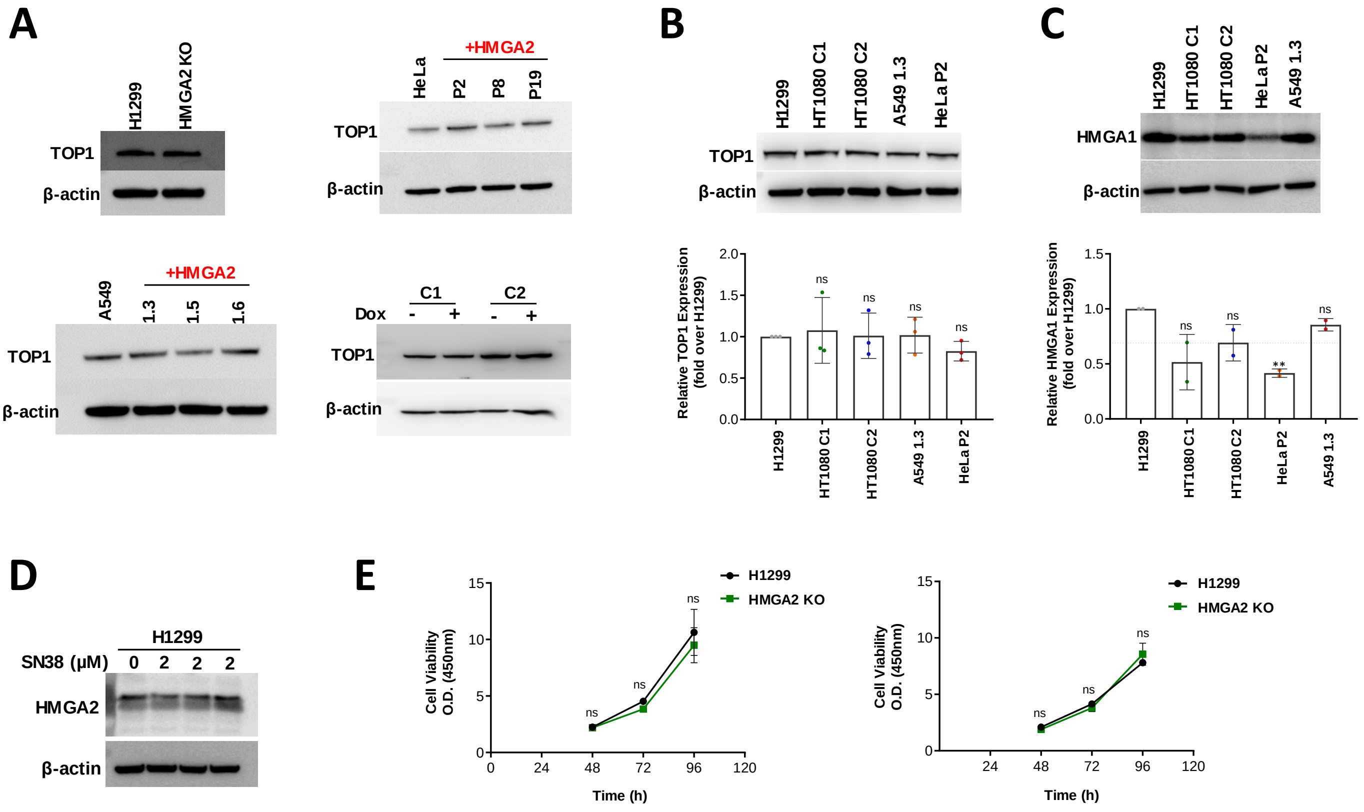

### S4 Fig

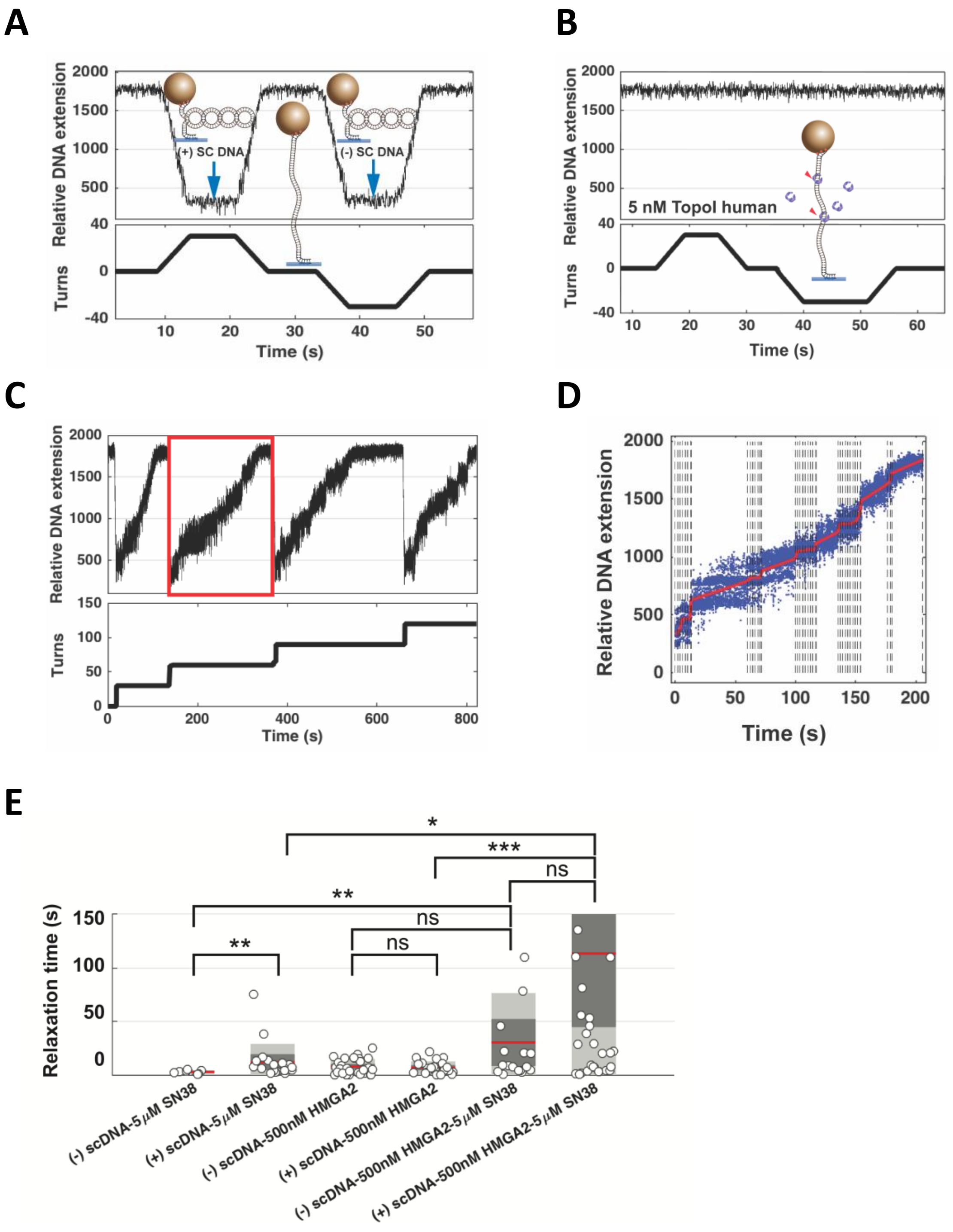

### S5 Fig

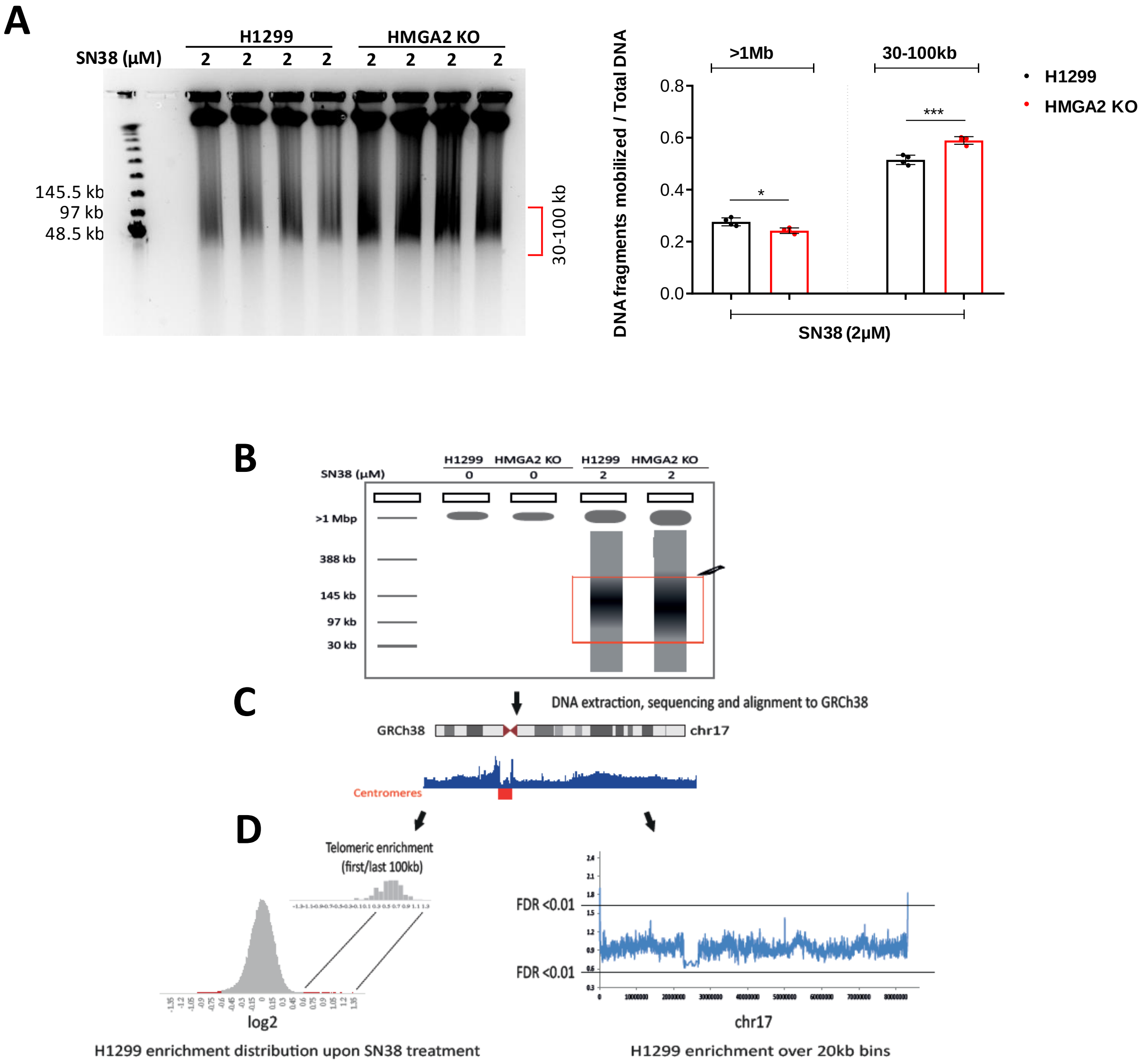

### S6 Fig

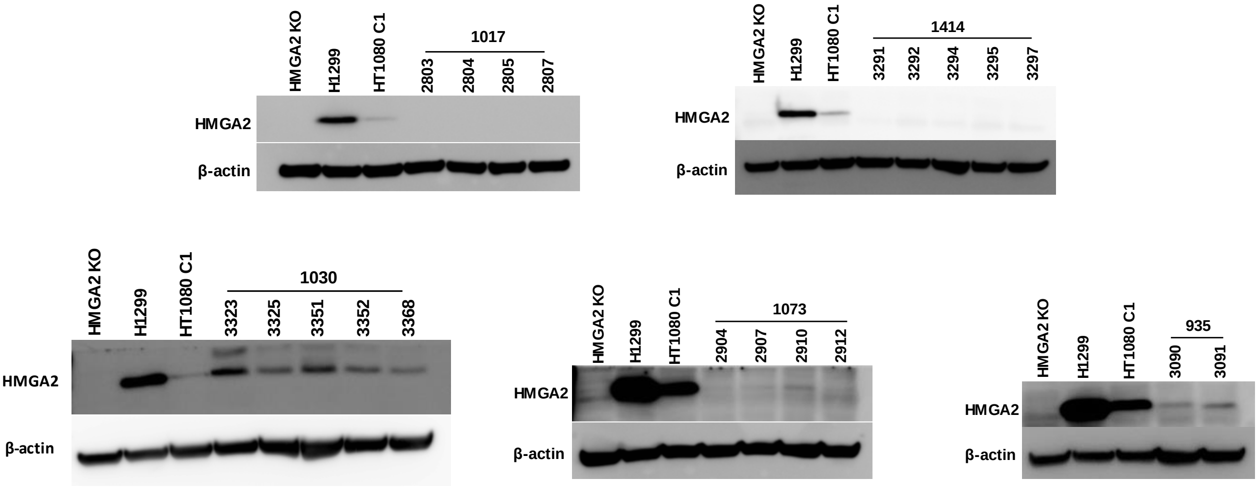
